# Evolutionary proteomics uncovers ciliary signaling components

**DOI:** 10.1101/153437

**Authors:** Monika Abedin Sigg, Tabea Menchen, Jeffery Johnson, Chanjae Lee, Semil P. Choksi, Galo Garcia, Henriette Busengdal, Gerard Dougherty, Petra Pennekamp, Claudius Werner, Fabian Rentzsch, Nevan Krogan, John B. Wallingford, Heymut Omran, Jeremy F. Reiter

**Affiliations:** Department of Biochemistry and Biophysics, Cardiovascular Research Institute, University of California, San Francisco, CA 94158, USA; Department of General Pediatrics, University of Muenster, Muenster 48149, Germany; Gladstone Institute of Cardiovascular Disease and Gladstone Institute of Virology and Immunology, San Francisco, CA 94158, USA; Department of Molecular Biosciences, Center for Systems and Synthetic Biology and Institute for Cellular and Molecular Biology, University of Texas at Austin, Austin 78712, Texas; Sars International Centre for Marine Molecular Biology, University of Bergen, Bergen 5008, Norway; Department of Cellular and Molecular Pharmacology, University of California, San Francisco, CA 94158, USA

## Abstract

Cilia are organelles specialized for movement and signaling. To infer when during animal evolution signaling pathways became associated with cilia, we characterized the proteomes of cilia from three organisms: sea urchins, sea anemones and choanoflagellates. From these ciliomes, we identified 437 high confidence ciliary candidate proteins conserved in mammals, including known regulators of Hh, GPCR and TRP channel signaling. The phylogenetic profiles of their ciliary association indicate that the Hh and GPCR pathways were linked to cilia before the origin of bilateria and TRP channels before the origin of animals. We demonstrated that some of the candidates not previously implicated in ciliary biology localized to cilia and further investigated ENKUR, a TRP channel-interacting protein that we identified in the cilia of all three organisms. In animals, ENKUR is expressed by cells with motile cilia, ENKUR localizes to cilia in diverse organisms and, in both *Xenopus laevis* and mice, ENKUR is required for patterning the left/right axis. Moreover, mutation of *ENKUR* causes situs inversus in humans. Thus, proteomic profiling of cilia from diverse eukaryotes defines a conserved ciliary proteome, reveals ancient connections to Hh, GPCR and TRP channel signaling, and uncovers a novel ciliary protein that controls vertebrate development and human disease.

## INTRODUCTION

The development of multicellular animals, involving complex cell movements, growth, patterning and differentiation, is orchestrated through intercellular communication. During and after development, some forms of cell-cell communications require cilia (Goetz and Anderson, 2010). Similarly, certain forms of intercellular communication that occur among single-celled organisms, such as mating in the unicellular green alga *Chlamydomonas*, involve cilia (Pan and Snell, 2000; Solter and Gibor, 1977; Wang et al., 2006). In addition to sensing signals from other cells, cilia in both unicellular protozoa and multicellular animals can sense environmental stimuli such as light and chemical cues such as odors (Bloodgood, 2010; Malicki and Johnson, 2016).

Cilia and the related structures, flagella, are comprised of microtubular cores called axonemes that emanate from basal bodies and are ensheathed by membranes that are compositionally distinct from the contiguous plasma membrane

(Carvalho-Santos et al., 2011; Satir and Christensen, 2007). In animals, some types of cilia are motile and others immotile. Vertebrates use non-motile primary cilia for detecting certain environmental stimuli and for intercellular communication, functions that require the localization of specific signal transduction machinery to cilia (Goetz and Anderson, 2010). Mammals use motile cilia and flagella for sperm locomotion and the generation of extracellular fluid flow (Satir and Christensen, 2007).

One critical role for motile cilia in many vertebrate embryos is in the specification of the left-right body axis. In mammals, motile cilia in the node, a cup-shaped structure on the ventral surface of the embryo, beat clockwise to generate a leftward flow that is required for left-right axis patterning (Marszalek et al., 1999; Nonaka et al., 1998; Supp et al., 1999). Motile cilia also clear lungs of mucous and debris, and circulate cerebrospinal fluid (Satir and Christensen, 2007).

Like primary cilia, motile cilia can also transduce signals required for animal development and adult tissue function. For example, cilia on human airway epithelial cells possess G-protein coupled receptors (GPCRs) that increase beat frequency in response to bitter compounds (Bloodgood, 2010; Shah et al., 2009). Cilia may also sense the leftward flow in left-right axis specification (McGrath et al., 2003).

Three signaling pathways associated with cilia are Hedgehog (Hh) signaling, some forms of GPCR signaling, and the activity of select Transient receptor potential (TRP) channels. The connections between cilia and Hh signaling, a critical regulator of embryonic development and adult homeostasis, are perhaps the best characterized, and many of the Hh signal transduction components localize to cilia (Goetz and Anderson, 2010). GPCRs that sense specific neurotransmitters and hormones also localize to neuronal primary cilia, suggesting that they have specific functions there (Hilgendorf et al., 2016). Similarly, TRP channel family members present in cilia regulate local calcium levels to modulate embryonic development (Delling et al., 2013).

Reflecting the diverse roles of cilia, ciliary defects cause a variety of human diseases with a wide range of manifestations, collectively called ciliopathies. Ciliopathy-associated phenotypes include retinal degeneration, polycystic kidney disease (PKD), skeletal malformations, bronchiectasis and left-right axis defects (Hildebrandt et al., 2011).

Because of the deep evolutionary conservation of cilia, the functions of cilia in single-celled organisms provide insights into the functions of animal cilia. For example, proteomic analysis of the *Chlamydomonas* flagellum and comparative genomics of ciliated and non-ciliated organisms has helped define the ciliary parts list and identify the genetic causes of ciliopathies (Avidor-Reiss et al., 2004; Li et al., 2004; Pazour et al., 2005).

Unlike the extensive conservation of ciliary structural proteins between *Chlamydomonas* and humans, many ciliary signaling proteins, including most components of the Hh pathway, are not conserved beyond animals (Adamska et al., 2007). We hypothesized that comparisons of the ciliary proteomes of animals and their close relatives would reveal new components of ciliary intercellular signaling pathways and help elucidate how signaling and cilia became associated over animal evolution. To test this hypothesis, we selected three organisms phylogenetically positioned to reveal ancestral ciliary functions at critical steps throughout animal evolution: sea urchins, sea anemones, and choanoflagellates. Sea urchins, early-branching non-chordate deuterostomes with embryonic cilia that are used for motility and Hh signaling (Warner et al., 2013), can provide insights into the signaling pathways associated with cilia before the emergence of chordates. Sea anemones are cnidarians (non-bilaterians) and like sea urchins, possess highly ciliated embryos. Sea anemones can reveal ciliary proteins that were present in early-branching animals, before bilaterians arose. As the closest known relative of animals, choanoflagellates are well positioned to shed light on the evolutionary origin of metazoans. These flagellated protozoa can elucidate ciliary proteins that evolved before the emergence of animals.

We identified ciliary proteins from each of these organisms by isolating their cilia and analyzing them by mass spectrometry. Comparison of the ciliary proteomes, which we refer to as evolutionary proteomics, revealed a core, shared ciliary proteome and helped illuminate the ancestry of ciliary signaling proteins, uncovering ancient links between cilia and GPCR, Hh and TRP channel signaling pathways. Evolutionary proteomics also identified previously uncharacterized, evolutionarily ancient ciliary proteins conserved in mammals, including a protein called ENKURIN (ENKUR). We found that ENKUR is highly conserved among ciliated eukaryotes. ENKUR orthologs from diverse animal species and choanoflagellates localize to cilia, suggesting a conserved ciliary function. Functional characterization of ENKUR revealed that it is expressed in tissues with motile cilia and is required for left-right axis patterning in *Xenopus* and mice, but is dispensable for ciliary motility in the mouse respiratory tract. Furthermore, mutations in human *ENKUR* may cause inherited situs inversus, revealing that evolutionary analysis of organellar proteomes can provide insight into the etiology of disease.

## RESULTS

### Ciliary proteomes of sea urchins, sea anemones and choanoflagellates

To identify ciliary proteins conserved between animals and a close relative of animals, we isolated cilia from the sea urchin *Strongylocentrotus purpuratus*, the sea anemone *Nematostella vectensis,* and the choanoflagellate *Salpingoeca rosetta* (Figure 1A). Sea urchins develop from triploblastic embryos that are covered with motile cilia (Figure 1B). Sea anemones develop from diploblastic embryos and, like sea urchins, have a highly ciliated larval stage. Choanoflagellate cells have a single, apical flagellum (Dayel et al., 2011).

**Figure 1.**
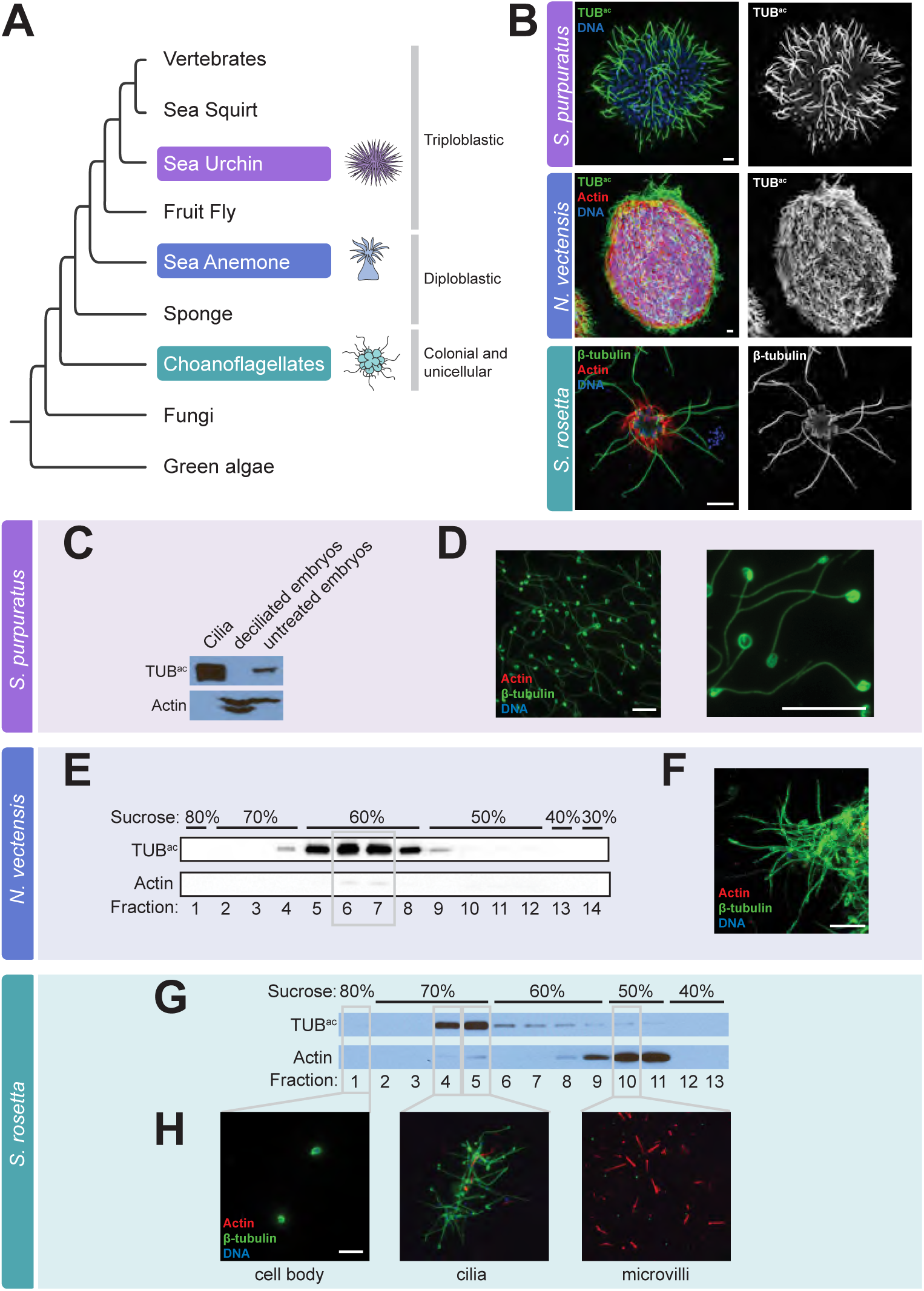
Isolation of cilia from sea urchins, sea anemones and choanoflagellates. **(A)** The phylogenetic relationship of the organisms studied in this work to other eukaryotes. **(B)** Immunofluorescent staining of cilia, marked by β-tubulin or Tubulin^ac^ (TUB^ac^) (green), in a sea urchin (*Strongylocentrotus purpuratus*) gastrula embryo, a sea anemone (*Nematostella vectensis*) planula larva, and a colony of choanoflagellate cells (*Salpingoeca rosetta*). Phalloidin staining of the sea anemone larva demonstrates the localization of Actin (red) to the cell bodies. Phalloidin staining of the choanoflagellate colony marks the collar of microvilli that surrounds each flagellum. Nuclei are stained with DAPI (blue). **(C)** Lysates of isolated *S. purpuratus* embryonic cilia, deciliated embryos, and intact embryos immunoblotted for TUB^ac^ and Actin. Cilia are enriched for TUB^ac^ and have undetectable amounts of Actin, whereas deciliated embryos have undetectable amounts of TUB^ac^. **(D)** Immunostaining of purified *S. purpuratus* cilia for b-tubulin (green), Actin (red) and nuclei (DAPI, blue) demonstrates that the axonemes of isolated cilia remain intact (β-tubulin) and confirms that no Actin or nuclei are detected in the ciliary fraction. **(E)** Immunoblot analysis of fractions from the sucrose step gradient purification of isolated *N. vectensis* cilia reveals that the 60% sucrose fractions contains TUB^ac^, peaking in fractions 6 and 7. **(F)** Immunostaining of fraction 7 for β-tubulin (green), Actin (red) and nuclei (DAPI, blue) confirms that the sea anemone ciliary fraction is replete with cilia (TUB^ac^) and contains little Actin and no detectable nuclei. **(G)** Immunoblot analysis of fractions from the sucrose step gradient purification of *S. rosetta* components reveal that TUB^ac^ is enriched in the 70% sucrose step and Actin is enriched in the 50% sucrose step. (**H**) Immunostaining of fractions for β-tubulin (green), Actin (red) and nuclei (DAPI, blue) confirms that cilia are enriched in the 70% sucrose step, cell bodies identified by DAPI staining are in the 80% sucrose step and microvilli marked by Actin are enriched in the 50% sucrose step. Scale bars, 5 μm for all images.

For each of these three species, all of which possess mono-ciliated cells, we isolated cilia from developmental stages selected to increase the likelihood of identifying ciliary signaling proteins relevant to development. As sea urchin gastrula stage cilia are required for development (Warner et al., 2013), we used a high salt shock to amputate cilia from gastrula stage sea urchin embryos (Figure 1B) and separate deciliated embryos from the isolated cilia by differential centrifugation (Auclair and Siegel, 1966; Stephens, 1986). This method resulted in efficient separation of whole cilia, including the axoneme and ciliary membrane, from the basal body and the rest of the cell (Stephens, 1995). Immunofluorescent imaging and immunoblot analysis demonstrated that the ciliary fraction was highly enriched for the ciliary components β-Tubulin and acetylated Tubulin (TUB^ac^) and Actin, a non-ciliary protein, was undetectable (Figure 1C and D). To enhance the detection of stage-specific ciliary signaling proteins, we isolated cilia from sea urchin embryos at both early and late gastrulation stages. To enrich for signaling proteins which are often less abundant than structural and motor proteins, we separated the axonemes and their associated proteins from the non-axonemal portion, also called the “membrane plus matrix” fraction (Figure 2A) (Pazour et al., 2005; Witman et al., 1972). SDS-PAGE revealed unique banding patterns for each fraction, indicating successful separation (Figure 2B).

**Figure 2.**
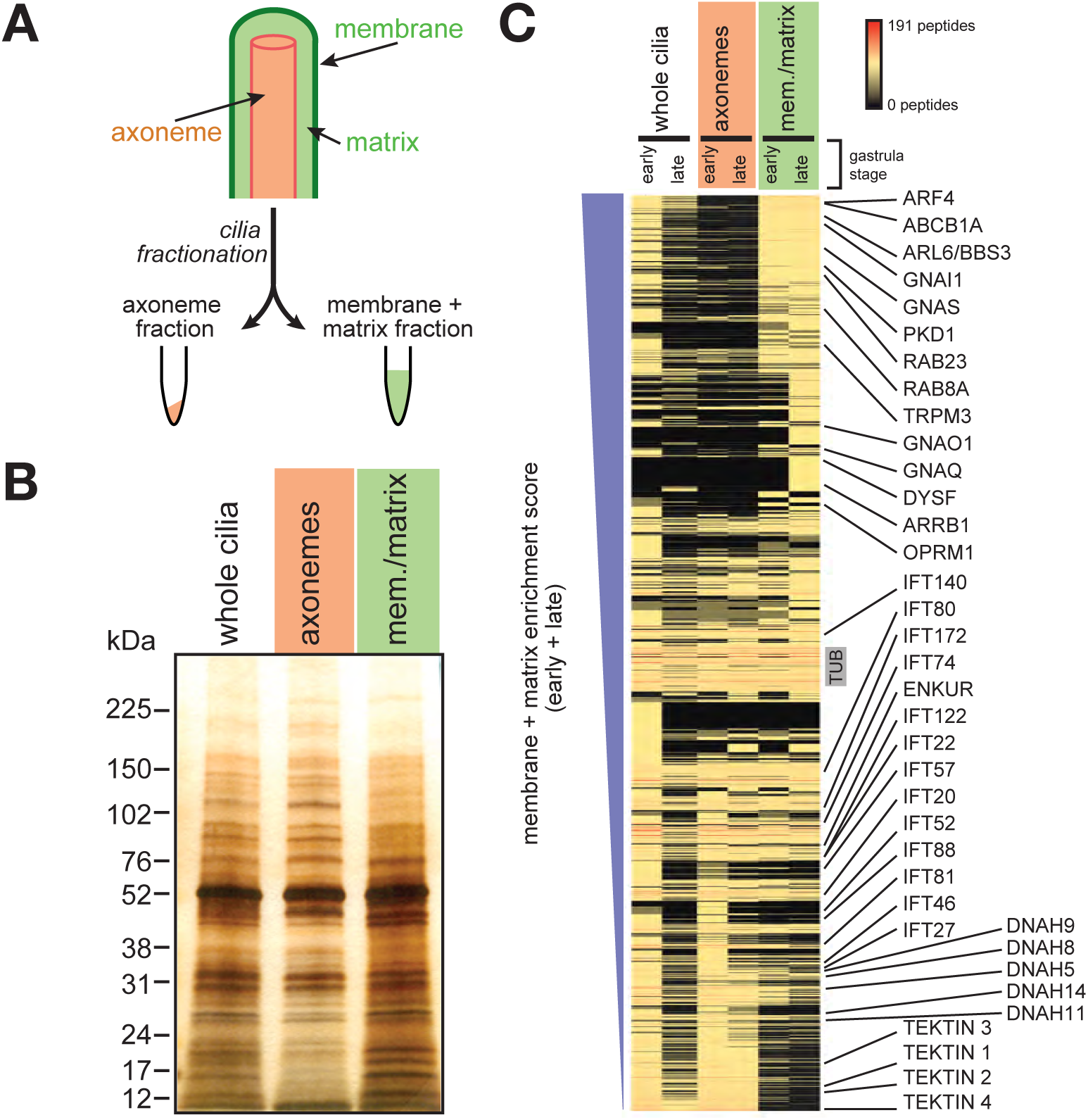
Characterization of the sea urchin ciliome. **(A)** A schematic of ciliary fractionation. Treatment of whole isolated *S. purpuratus* cilia with detergent followed by centrifugation separates cilia into two fractions, axonemes and membrane + matrix. **(B)** Silver stain of isolated *S. purpuratus* whole cilia, axonemes and membrane + matrix (mem./matrix) fractions resolved by SDS-PAGE. The protein banding pattern of the axoneme fraction differs from that of the membrane + matrix fraction. **(C)** Proteins sorted by their membrane + matrix enrichment score (see Methods). The unique peptide count for each protein is represented in the heat map. Known axonemal proteins (*e.g.*, IFT components, axonemal Dyneins and Tektins) are enriched in the axonemal fraction. Known ciliary membrane and membrane-associated proteins (*e.g.*, PKD1, ARF4, RAB8A) are enriched in the membrane + matrix fraction.

To isolate sea anemone cilia, we developed a deciliation protocol based on high salt shock. After separating cilia from sea anemone larvae of the planula stage (the stage that follows gastrulation) (Figure 1B), we purified the cilia by sucrose gradient centrifugation and fractionation. Fractions high in TUB^ac^ contained little Actin and were comprised of intact cilia (Figure 1E and F).

We discovered that dibucaine, a compound used to isolate flagella from *Chlamydomonas* and *Tetrahymena,* caused choanoflagellate cells in rosette colonies to release their flagella (Thompson et al., 1974; Witman, 1986). In *Chlamydomonas,* diabucaine severs cilia distal to the basal body and transition zone (Sanders and Salisbury, 1994). After amputating flagella from colonial choanoflagellates (Figure 1B), we used sucrose gradient centrifugation and fractionation to separate microvilli and cell bodies from isolated cilia. Again, immunofluorescent imaging and immunoblot analysis revealed that the ciliary fraction was highly enriched for TUB^ac^, whereas the microvilli fraction was enriched for Actin (Figure 1G and H).

To identify the ciliary proteomes of these species, we analyzed the isolated cilia and flagella by mass spectrometry. We performed mass spectrometric protein profiling on unfractionated purified sea urchin cilia (whole cilia), the isolated axonemal fraction (axonemes) and the non-axonemal fraction (membrane plus matrix). Due to the tractability of obtaining large quantities of sea urchin cilia, we were able analyze cilia isolated from three separate embryo cultures at two different developmental stages. We separated the peptides from unfractionated whole cilia using two distinct methods of chromatography: pre-fractionated by hydrophilic interaction chromatography followed by C18 LC-MS/MS or LC-MS/MS using a high resolution C18 column and an extended reverse phase gradient. For the mass spectrometry data analyses, we chose a low-stringency cut-off to capture signaling proteins: proteins with a unique peptide count of 2 or more were included for future analysis (Table S1).

For isolated sea anemone cilia, we analyzed whole cilia from sucrose gradient fractions containing peak levels of TUB^ac^ from two individual embryo cultures (Figure 1E, fractions 6 and 7). As with the sea urchin, to further increase the diversity of peptides that we could identify, we separated peptides using the two distinct methods of chromatography described above.

To obtain a sufficient amount of material for mass spectrometry of choanoflagellate cilia, we performed two separate cilia preparations and pooled the samples before analysis. The sample was fractionated by high resolution reverse phase before mass spectrometry. To identify proteins selectively enriched in choanoflagellate flagella, we analyzed fractions containing cell bodies (fraction 1) and microvilli (fraction 10) alongside the ciliary fractions (fractions 4 and 5) (Figure 1G). We defined proteins with a ciliary fraction peptide count higher than that of the cell body and microvilli fraction as candidate members of the choanoflagellate ciliary proteome, or ‘ciliome′.

Metazoan developmental signaling pathways are proposed to have emerged with the advent of multicellularity or with major transitions in animal evolution (Pincus et al., 2008; Pires-daSilva and Sommer, 2003). Many signal transduction components are membrane-associated, suggesting that the membrane plus matrix fraction of sea urchin cilia would be enriched for signal transduction components relative to the axonemal fraction. Therefore, we calculated the degree of membrane plus matrix enrichment for each sea urchin protein by comparing its peptide count in the membrane plus matrix fraction to that of the axonemal fraction (Figure 2C and Table S2).

Comparing the data from the sea urchin whole cilia, axonemes and membrane plus matrix revealed that some proteins identified in ciliary fractions were not detected in the whole cilia data (Figure 2C), suggesting that fractionation increased the sensitivity of peptide detection. Conversely, a small subset of proteins detected in the early gastrula whole cilia fraction were not detected in the ciliary fraction, likely due to enrichment by hydrophilic interaction chromatography. All other sea urchin samples were analyzed exclusively by reverse phase chromatography. As expected, previously identified axonemal proteins (e.g., IFTs, motor proteins, Tektins) were enriched in the axonemal fraction (Figure 2C). Also as expected, previously identified ciliary membrane-associated proteins (e.g., PKD1, BBS3, RAB8A) were enriched in the membrane plus matrix fraction (Figure 2C). Moreover, membrane proteins not previously associated with cilia were enriched in the membrane plus matrix, including Dysferlin (DYSF), ATP-binding cassette, sub-family B (MDR/TAP), member 1A (ABCB1A), Transient receptor potential cation channel subfamily M member 3 (TRPM3).

### Definition of a conserved ciliome identifies novel ciliary proteins

From the isolated cilia of three organisms, we identified peptides corresponding to over 3,000 proteins in total (Table 1 and S1). As we are most interested in the evolution of ciliary signaling in the progression from the unicellular ancestors of animals to humans, we focused on those proteins with a homolog in mammals. BLAST identified 1,266 ciliome proteins with a homolog in mouse (Table 1 and S1), a set we termed the ‘conserved ciliome′. As expected based on their evolutionary distances to mammals, the sea urchin ciliome contains the greatest number of mammalian homologs (1,012), followed by the sea anemone (511) and choanoflagellate ciliomes (311), respectively (Table 1, Figure 3A).

**Table 1.** Mass spectrometry data.

|  | Choanoflagellate | Sea anemone | Sea urchin | Total |
| --- | --- | --- | --- | --- |
| number of proteins identified | 464<sup>a</sup> | 991<sup>a</sup> | 2,222<sup>a</sup> | n/a |
| proteins with homolog in mouse | 311 | 511 | 1,012 | 1,266 |
| *Known cilia proteins* |  |  |  |  |
| SYSCILIA Gold Standard genes | 41% | 55% | 69% | n/a |
| published cilia data sets | 208 | 360 | 644 | 765 |
| disease genes | 67 | 93 | 212 | 241 |
| ciliopathy genes | 36 | 47 | 69 | 73 |

<sup>a</sup> multiple protein isoforms encoded by single genes are present in these data

**Figure 3.**
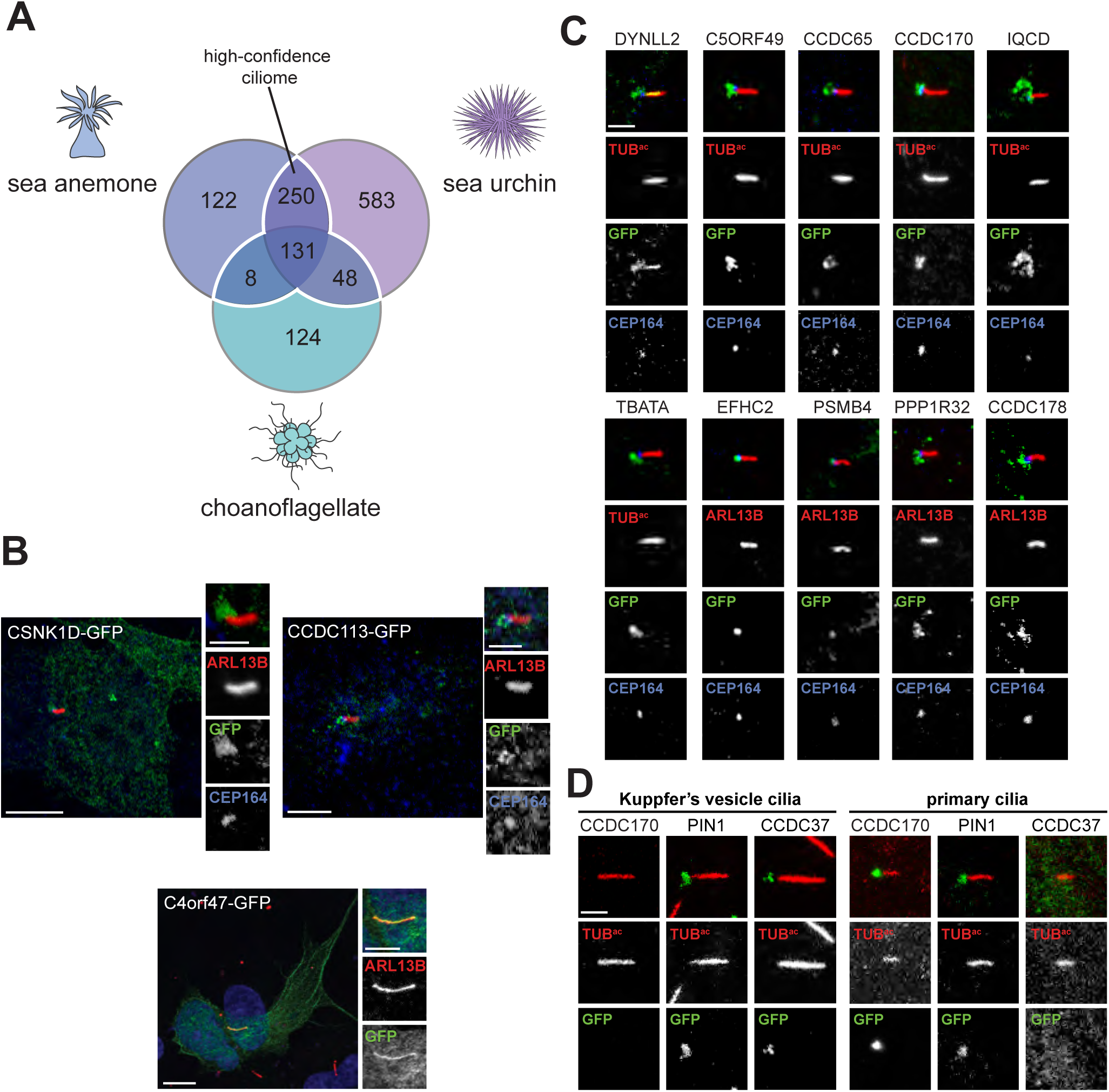
Defining an evolutionarily conserved ciliome. **(A)** Venn diagram of the overlap of *S. purpuratus*, *N. vectensis* and *S. rosetta* ciliomes. The white line encompasses the high-confidence ciliary proteins present in 2 or more ciliomes. Only proteins that possess a mouse homolog (BLAST E value ≤ 1e-5) are included. **(B)** Immunofluorescent staining for cilia (ARL13B, red) and candidate proteins fused to GFP (green) expressed in IMCD3 cells. Human C4orf47-GFP localizes to cilia and cytoplasmic microtubules. Fusions of human CSNK1D and CCDC113 with GFP predominantly co-localize with the basal body component CEP164 (blue). For C4orf46-GFP, nuclei are stained with Hoechst (blue). Scale bar for whole cell images, 5 μm. Scale bar for cilia only images, 2.5 μm. **(C)** Ten GFP-tagged human proteins, out of 49 randomly selected proteins from the high-confidence ciliome, localize to cilia or the ciliary base of IMCD3 cells. Immunofluorescent staining marks GFP-tagged proteins (green), cilia are indicated by ARL13B or TUB^ac^ (red), as indicated, and the basal body is highlighted by CEP164 (blue). Scale bar, 2.5 μm. **(D)** Immunofluorescent staining of human proteins fused to GFP (green) in the *D. rerio* embryo (somite stage 6-10). Cilia are marked by staining for TUB^ac^ (red). Cilia from within the Kuppfer’s vesicle and primary cilia found outside the Kuppfer’s vesicle are depicted. Scale bar, 2.5 μm.

As a first assessment of the ciliome, we compared the ciliary members of the SYSCILIA Gold Standard database, a list of well characterized proteins involved in cilia biology (van Dam et al., 2013), to each organism’s ciliome. Of the SYSCILIA Gold Standard proteins present in cilia, 69% were present in the sea urchin ciliome (Tables 1 and S3). The sea anemone ciliome contained 55% of the SYSCILIA Gold Standard ciliary proteins and the choanoflagellate ciliome contained 41%. Previous ciliary proteomics or genomics studies identified less than half the number of the ciliary SYSCILIA Gold Standard members, suggesting that the ciliomes encompasses a large proportion of ciliary components (Avidor-Reiss et al., 2004; Ishikawa et al., 2012; Li et al., 2004; Mick et al., 2015; Ostrowski et al., 2002; Pazour et al., 2005).

437 mouse homologs were represented in at least two of the ciliomes, providing confidence that they represent bona fide ciliary components (Figure 3A, Table S1 highlighted in blue). To begin to assess whether this ‘high-confidence ciliome’ encompasses evolutionarily conserved ciliary components, we compared it to proteins implicated in ciliary biology by 19 genomic, expression and proteomic studies in diverse unicellular and multicellular ciliated organisms (see Table S4 for list of studies and associated organisms) (Avidor-Reiss et al., 2004; Blacque et al., 2005; Boesger et al., 2009; Broadhead et al., 2006; Cao et al., 2006; Chen et al., 2006; Choksi et al., 2014; Efimenko et al., 2005; Ishikawa et al., 2012; Laurençon et al., 2007; Li et al., 2004; Liu et al., 2007; Mayer et al., 2008; Merchant et al., 2007; Mick et al., 2015; Narita et al., 2012; Ostrowski et al., 2002; Pazour et al., 2005; Smith et al., 2005). 76% of the high-confidence ciliome proteins has been previously implicated in ciliary biology (Table S1).

As the closest known living relative of animals, we hypothesized that choanoflagellate cilia would contain proteins conserved in mammals that are not present in the cilia of protozoa more distantly related to animals. To test this hypothesis, we compared the choanoflagellate ciliome to that of *Chlamydomonas reinhardtii,* the flagella of which have been well characterized by mass spectrometry (Pazour et al., 2005). We found that out of the 465 proteins in the choanoflagellate ciliome, 267 did not have a homolog in the *C. reinhardtii* ciliome (Table S3). Of the choanoflagellate ciliome proteins not present in the *C. reinhardtii* ciliome, over half (157 proteins) had a homolog in the mouse, indicating that these protein are not unique to choanoflagellates and may represent ciliary proteins that emerged after the evolutionary divergence of *Chlamydomonas* from the lineage that would give rise to animals.

As a final test of whether the ciliomes represent a broad survey of ciliary components, we examined whether ciliopathy-associated proteins were identified in the conserved ciliome. Seventy-four of the conserved ciliome proteins are mutated in a ciliopathy, including 11 proteins associated with Bardet-Biedl-Syndrome (BBS), both proteins associated with autosomal recessive PKD, 13 proteins associated with short rib thoracic dysplasia, and 21 proteins associated with primary ciliary dyskinesia (PCD) (Table S1).

Thus, we hypothesized that high-confidence ciliome constituents not associated with ciliopathies are candidates that may explain the etiologies of orphan ciliopathies. For example, one high-confidence ciliome member, coiled-coil domain containing 63 (CCDC63), was recently identified as being essential for mouse sperm flagella formation, suggesting that it could be a ciliary protein linked to male fertility (Young et al., 2015). Other ciliome constituents may also indicate ciliary etiologies for syndromes not previously recognized as ciliopathies. For example, WD-repeat domain 65 (WDR65, also known as CFAP57), was identified in all three ciliomes and is associated with van der Woude syndrome, a craniofacial malformation with features similar to that of Orofaciodigital syndrome, a recognized ciliopathy (Rorick et al., 2011).

We also assessed whether these ciliomes have predictive power to identify previously undescribed ciliary components. C4orf22 (1700007G11Rik), C9orf116 (1700007K13Rik), C10orf107 (1700040L02Rik), MORN repeat containing 5 (MORN5), IQ motif containing D (IQCD), armadillo repeat containing 3 (ARMC3), UBX domain protein 11 (UBXN11), MYCBP associated protein (MYCBPAP), coiled-coil domain containing 96 (CCDC96), IQ motif and ubiquitin domain containing (IQUB), EF hand calcium binding domain 1 (EFCAB1), DPY30 domain containing 1 (DYDC1) all prominently localize to the multicilia of fallopian tube cells, suggesting that they may be previously unrecognized components of motile cilia (Uhlén et al., 2015). We expressed LAP-tagged versions of select candidates and assessed their localization in IMCD3 cells. C4orf47 (1700029J07Rik), for example, localizes to cytoplasmic microtubules and cilia (Figure 3B). Others, including Coiled-coil domain containing 113 (CCDC113) and Casein kinase 1δ (CSNK1D) localized predominantly to the base of cilia (Figure 3B).

Many proteins in the high-confidence ciliome are known to localize to and function within mammalian cilia, but others, like C4orf47, have not previously been identified as ciliary proteins. To estimate what proportion of proteins in the high-confidence ciliome are bona fide ciliary proteins in mammals, we randomly selected 49 proteins from the high-confidence ciliome and tested if the GFP-tagged human homolog localized to cilia of IMCD3 cells (Table S5). We also tested the localization 30 randomly selected human proteins not identified in the ciliome. Based on proportion of the human genome encoding known ciliary proteins, we expected that 1-3 proteins would localized to cilia. Four of the randomly selected proteins localized to cilia (Figure S1A), suggesting that the test is specific. Of the 49 randomly selected candidate ciliary proteins, 10 localized to the cilium or the ciliary base (Figure 3C and S1B). Included in the 49 proteins were 8 SYSCILIA Gold Standard proteins, one of which localized to cilia, indicating that this test is not sensitive. Based on the proportion of candidate ciliary proteins that localized to cilia and the low sensitivity of the localization assay, greater than 20% of the proteins in the high-confidence ciliome are likely to localize to human cilia.

We hypothesized that some proteins in the ciliome localize specifically to motile cilia, and not to primary cilia. To test this hypothesis, we determined whether proteins that did not localize to IMCD3 primary cilia can localize to the motile cilia of the zebrafish Kupffer’s vesicle. CCDC37 (CFAP100) and PIN1, two proteins that did not localize to primary cilia in IMCD3 cells, localized to motile cilia of the zebrafish embryo (Figure 3D and S1C). Unlike CCDC37, PIN1 localized to some, but not all primary cilia in the zebrafish embryo. Conversely, CCDC170 localized to the base of primary cilia in IMCD3 cells and in the zebrafish embryo, but did not localize to the motile cilia of Kupffer’s vesicle. These data reveal that some proteins localize specifically to either motile or primary cilia.

### Comparison of evolutionarily diverse ciliomes identifies signaling proteins

Many ciliary components show a taxonomic distribution consistent with being present in the last common eukaryotic ancestor and lost from organisms that have lost cilia, such as many plants and fungi (Li et al., 2004). To test whether components of the sea urchin, sea anemone and choanoflagellate ciliomes display a taxonomic distribution paralleling the distribution of cilia, we used a computational algorithm, CLustering by Inferred Models of Evolution (CLIME), to map proteins onto a phylogeny that includes prokaryotes and 136 ciliated and unciliated eukaryotes (Li et al., 2014). 15% of ciliome components displayed a taxonomic distribution closely mirroring the presence of cilia (Figure 4A, S2 and Table S6). In addition to many recognized ciliary components, such as IFT and BBSome, high-confidence ciliome components, such as ENKUR, TRPM3, Zinc finger MYND-type containing 12 (ZMYND12) and Sperm tail PG rich repeat containing 2 (STPG2) showed distributions consistent with co-evolving with cilia. The presence of these genes in the genomes of diverse ciliated eukaryotes suggests that they may have evolutionarily ancient functions in cilia.

**Figure 4.**
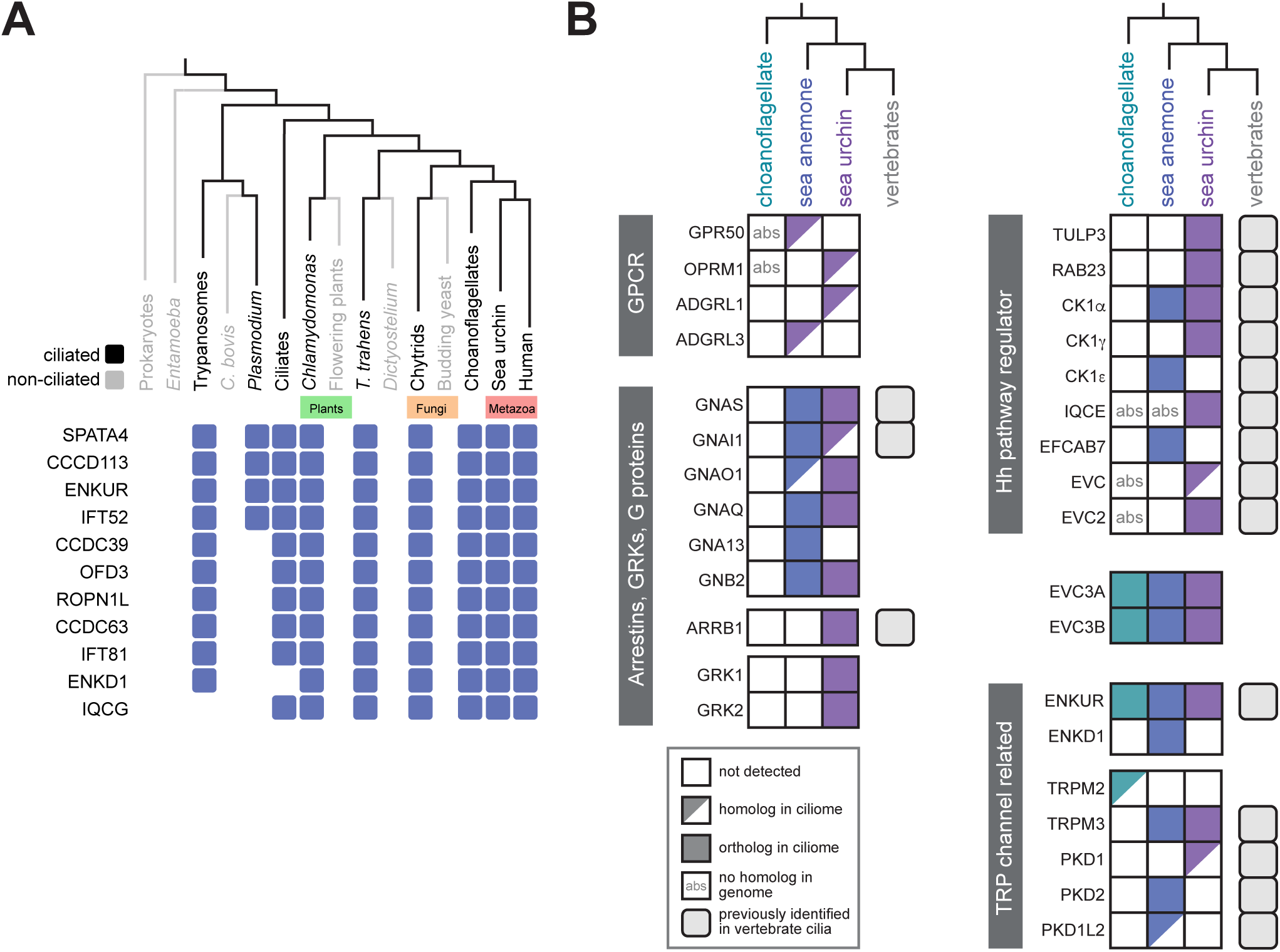
The distribution of signal transduction components detected in ciliomes. **(A)** Select ciliome proteins display a phylogenetic profile closely reflecting the distribution of cilia as based on CLIME analysis. Ciliated organisms are shown in black and non-ciliated organisms are shown in grey. A blue box indicates that a homolog is present in the organism or group of organisms represented in the phylogeny. For the complete list of ciliome proteins with distributions overlapping with ciliation within a phylogeny of 136 organisms, see Figure S2. For clades represented by more than one organism (*i.e.* Choanoflagellates, Budding yeast, Chytrids, Flowering plants, Ciliates, *Plasmodium*, Trypanosomes, *Entamoeba* and Prokaryotes) a protein was considered to be present in the clade if one or more organisms possessed the protein. **(B)** The ciliome of each organism contains diverse signal transduction components, most of which have homologs in vertebrates and some of which have not been previously associated with cilia. Proteins were considered orthologs if they were reciprocal best BLAST hits. The top BLAST hit in *M. musculus* was considered a homolog of the *S. purpuratus*, *N. vectensis* and *S. rosetta* query protein if the two proteins were not reciprocal BLAST matches (E value ≤ 1e-5).

To trace the ancestry of known ciliary signaling pathways, we also examined whether previously identified components of these pathways were present in the sea urchin, sea anemone and choanoflagellate ciliomes. Choanoflagellate genomes do not contain genes for most Hh pathway components, suggesting that they do not use Hh signaling for intercellular communication (King et al., 2008). Analysis of whether ciliary Hh pathway components were present in the sea urchin and sea anemone ciliomes revealed that, although canonical Hh pathway proteins such Patched1, Smoothened, and the GLIs were not detected, both positive regulators of ciliary Hh signaling, such as Casein Kinase 1 isoform γ (CK1γ) (Li et al., 2016) and negative regulators of ciliary Hh signaling, such as TULP3 (Chávez et al., 2015; Garcia-Gonzalo et al., 2015; Li et al., 2016; Mukhopadhyay et al., 2013) were detected (Figure 4B). Similarly, we detected Ellis van Creveld protein 2 (EVC2) and a homolog of EVC in the sea urchin ciliome. In mammals, EVC and EVC2 localize to the proximal cilium and act as tissue-specific positive regulators of the Hh pathway (Caparrós-Martín et al., 2013; Dorn et al., 2012; Yang et al., 2012). Mutations in the genes that encode these proteins cause Ellis–van Creveld syndrome and Weyers acrofacial dysostosis, ciliopathies characterized by skeletal and craniofacial defects and polydactyly (Ruiz-Perez et al., 2000; Ye et al., 2006). Two EVC and EVC2 interacting proteins, IQ domain-containing protein E (IQCE) and EF-hand calcium binding domain-containing protein 7 (EFCAB7), regulate Hh signaling by anchoring EVC and EVC2 to the proximal cilium (Pusapati et al., 2014). IQCE was present in the sea urchin ciliome and EFCAB7 was present in the sea anemone ciliome.

In addition to EVC and EVC2, we identified two paralogs, EVC3.1 and EVC3.2 (Pusapati et al., 2014), in the choanoflagellate, sea anenome and sea urchin ciliomes (Figure 4B). The presence of EVC3 in the ciliomes suggests that the association between cilia and the EVC family extends beyond metazoans to early-branching eukaryotes. As EVC3.1 and EVC3.2 are missing from the genomes of some mammals, including humans, it appears that, whereas mammals have preserved much of the ciliary repertoire of basal organisms, some have selectively lost at least two ciliary proteins.

Several of the largest class of receptors, GPCRs, localize to cilia. For example, odorant receptors act at olfactory epithelia cilia (Goetz and Anderson, 2010). We identified four GPCRs that have mouse homologs, none of which have been previously reported in cilia (Figure 4B). Other members of the GPCR signaling pathway, including G proteins, Arrestins and GPCR kinases (GRKs), were identified in both the sea urchin and sea anemone ciliomes, suggesting that not just GPCRs, but GPCR signal transduction has an ancient evolutionary connection to cilia that precedes the advent of bilateria.

To assess whether some of the identified GPCR signaling proteins can localize to mammalian cilia, we expressed a GFP-tagged version of a sea urchin homolog of Opioid receptor μ1 (OPRM1) that we named OPRM1L in RPE-1 cells and found that it localized to cilia (Figure S3A). The GPCR signal transduction component β-Arrestin-1/2 can localize to one end of cilia (Pal et al., 2016). As b-Arrestin-1 (ARRB1) was detected in the sea urchin ciliome, we also examined the localization of sea urchin ARRB1-GFP in RPE-1 cells and found that, like the mammalian ortholog, it was enriched at one ciliary end (Figure S3B).

Unlike Hh and GPCR signaling proteins, TRP channels and TRP channel-associated protein were detected in all three ciliomes (Figure 4B). PKD family members, previously known to localize to cilia, were present in sea anemone and sea urchin cilia, as was TRPM3, a channel that has multiple proposed functions, including heat sensation (Held et al., 2015). A homolog of TRPM2 was identified in the choanoflagellate cilia, suggesting that the association of TRP channels with cilia arose before the emergence of animals. Also, Enkurin (ENKUR), a TRP channel interacting protein (Sutton et al., 2004), was detected in the ciliomes of all three organisms and the Enkurin domain-containing protein, ENKD1, was detected in the sea anemone ciliome. ENKUR was originally identified as a TRP channel interactor expressed by sperm (Sutton et al., 2004). The identification of ENKUR orthologs in choanoflagellate, sea anemone and sea urchin ciliomes suggests a conserved ciliary function. As evolutionary conservation is associated with increased likelihood of affecting fitness and disease (Hirsh and Fraser, 2001; Wan et al., 2015), we hypothesized that ENKUR may have important biological functions in animals.

### ENKUR is a ciliary protein critical for left-right axis specification in *Xenopus* and mice

From the analysis of cilia of diverse organisms, we sought to uncover conserved proteins whose functions in ciliary biology had been unappreciated. As mammalian ENKUR has been detected at the sperm flagellum (Sutton et al., 2004), we wondered if ENKUR had functions in cilia and if those functions were conserved in earlier branching animals. In addition to identifying ENKUR in the ciliomes of choanoflagellates, sea anemones and sea urchins we found *Enkur* in the genome of diverse ciliated eukaryotes suggesting that it may have an evolutionarily ancient function in cilia (Figure S4).

To determine where in the invertebrate embryo *Enkur* is expressed we used in situ hybridization to identify *Enkur* expression in sea anemone and sea urchin embryos throughout gastrulation. In sea anemone embryos, early in gastrulation, *Enkur* was expressed in most cells, with higher expression in a subset of cells distributed throughout the embryo (Figure 5A and S5A). Midway through gastrulation and into the late gastrula stage, *Enkur* expression was maintained in the ectoderm and became enriched at the aboral end of the sea anemone embryo, coincident with the apical organ, a group of cells that bear long cilia (termed the “ciliary tuft”) and is hypothesized to have sensory function (Rentzsch et al., 2008).

**Figure 5.**
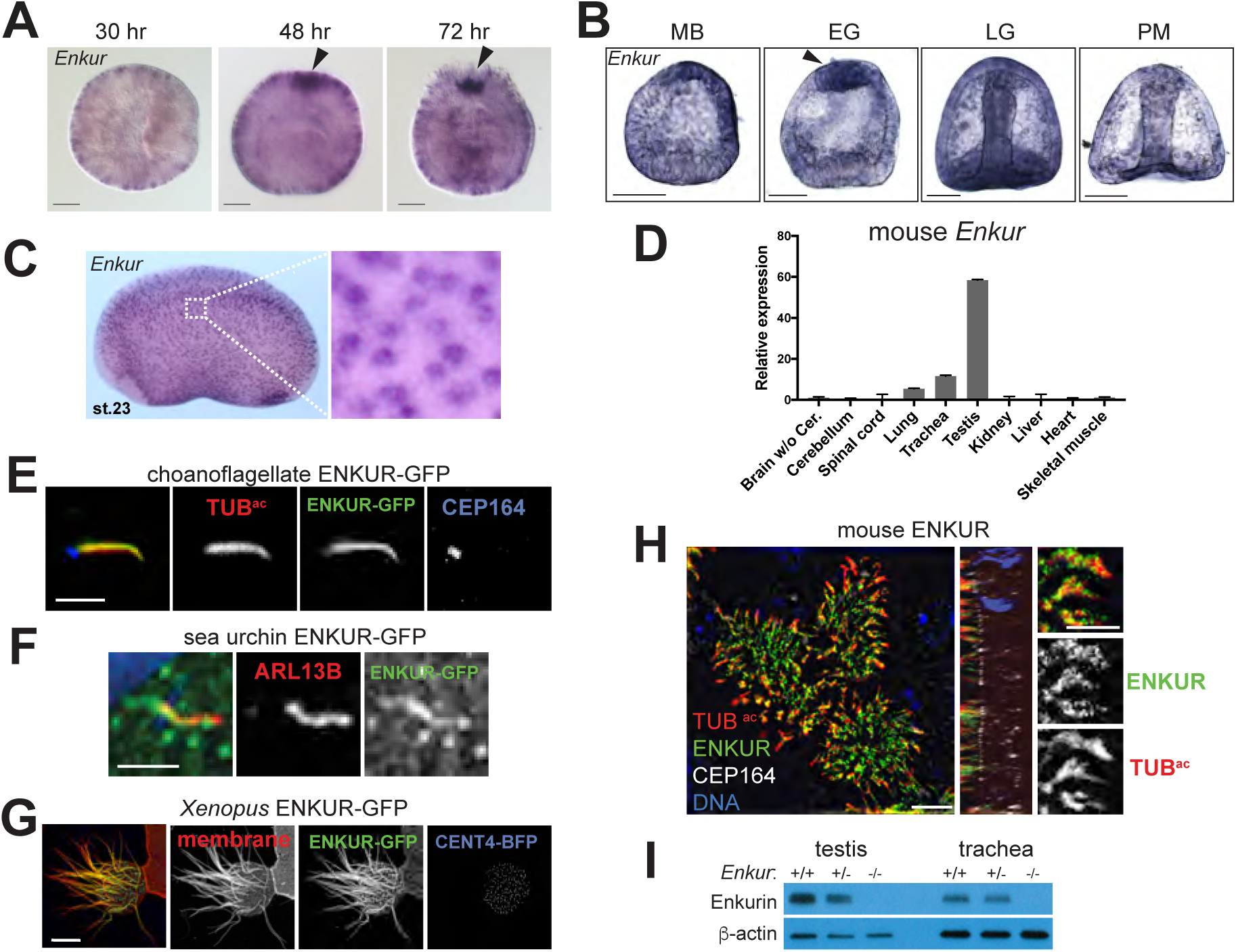
ENKUR is a conserved ciliary protein expressed by cells with motile cilia. (**A**) Whole mount in situ hybridization for *Enkur* in *N. vectensis* embryos at various developmental stages. *Enkur* is expressed throughout the embryo and is enriched at the aboral pole (arrowhead) in 48 and 74 hour embryos. Scale bar, 50 μm. (**B**) In situ hybridization for *Enkur* in *S. purpuratus* embryos at mesenchyme blastula (MB), early gastrula (EG), late gastrula (LG) and prism (PM) stages. Scale bar, 50 μm. *Enku*r is expressed in all cells and enriched at the apical pole in EG embryos. (**C**) In situ hybridization of stage 23 *X. laveis* embryo shows expression of *Enkur* in epidermal cells. (**D**) qRT-PCR measurement of *Enkur* expression in isolated adult mouse lungs, trachea and testis. Error bars represent SDs from 6 technical replicates. Expression was validated using 2 distinct primer pairs. **(E)** Immunofluorescent staining of *S. rosetta* ENKUR fused to GFP (green), cilia (TUB^ac^, red) and the basal body (CEP164, blue) expressed in IMCD3 cells. Scale bar, 2.5 μm. **(F)** Immunofluorescent staining of cilia (ARL13B, red) and a fusion of *S. purpuratus* ENKUR with GFP (green) expressed in RPE-1 cells. ENKUR-GFP localizes to cilia. Nuclei are stained with Hoechst (blue). Scale bar, 2.5 μm. **(G)** A multi-ciliated epidermal cell of a stage 23 *X. laevis* embryo expressing *X. laevis* ENKUR fused to GFP (green), membrane-red fluorescent protein (red), marking the plasma and ciliary membranes, and Centrin 4 (CENT4)-blue fluorescent protein (BFP, blue) to mark the basal bodies. ENKUR localizes along the length of cilia. Scale bar, 10 μm. (**H**) Immunofluorescent staining of primary cultured mouse tracheal epithelial cells for ENKUR (green), cilia (TUB^ac^, red), and the basal body (CEP164, white), and imaged using structured illumination microscopy. ENKUR localizes to mouse tracheal epithelial cilia. Scale bar for whole cells (top and center panels), 5 μm. Scale bar for cilia (bottom panel), 2.5 μm. (**I**) Immunoblotting for ENKUR protein in testis and trachea lysates from littermate control and *Enkur*^-/-^ mice. ENKUR is expressed in testis and trachea, whereas *Enkur*^-/-^ mice do not produce detectable ENKUR.

In the sea urchin embryo, *Enkur* is expressed in all cells from the mesenchyme blastula stage, throughout gastrulation and into the prism stage, with peak expression during gastrulation. Strikingly, in early gastrulation, *Enkur* expression is enriched at the apical end of the embryo where the ciliary tuft is located (Figure 5B and S5B). The expression pattern of *Enkur* in sea urchin and sea anemone embryos suggests a conserved function for ENKUR in the ciliary tuft that extends from early-branching eumazoans (all animals except sponges and placozoans) to dueterostomes.

To determine if, as in the sea urchin and sea anemone, *Enkur* is expressed in motile ciliated cells in vertebrate embryos, we used in situ hybridization to examine its expression in *Xenopus laevis* embryos. We found that *Enkur* is expressed in motile ciliated cells distributed throughout the epidermis (Figure 5C). In mouse, we confirmed by quantitative PCR that *Enkur* is expressed exclusively in mouse tissues that possess motile cilia, such as the testes and trachea (Figure 5D and (Sutton et al., 2004)).

To begin to assess the subcellular localization of sea urchin, sea anemone and choanoflagellate ENKUR, we GFP-tagged ENKUR and analyzed its localization in ciliated mammalian cells. We found that GFP-tagged sea urchin and choanoflagellate ENKUR proteins localized to cilia if expressed in mammalian cells (Figure 5E and F). To determine if ENKUR localizes to cilia in vertebrates, we expressed GFP-ENKUR in *X. laevis* embryos and found that it localized to the epidermal cell motile cilia (Figure 5G). Similarly, mouse ENKUR localized to the motile cilia of tracheal epithelial cells (Figure 5H and S5C) and the mammalian ortholog of an ENKUR-related protein, ENKD1, localized predominantly to the base of cilia (Figure S5D). To further analyze ENKUR expression in the mouse, we utilized *Enkur*^-/-^ mice. Western blot corroborated the ENKUR immunofluorescent staining, indicating that the protein is expressed in trachea, and is absent from the trachea of *Enkur*^-/-^ mice (Figure 5I).

In mammals, specification of the left-right axis requires both ciliary motility and PKD2-dependent ciliary signaling (McGrath et al., 2003; Nonaka et al., 1998; Pennekamp et al., 2002; Yoshiba et al., 2012). PKD2, a member of the TRP family of ion channel proteins, forms a ciliary complex with PKD1. A homologs of PKD1 was detected in the ciliome of sea urchin and an ortholog of PKD2 was detected in the ciliome of sea anenomes (Figure 4B). The localization of ENKUR to motile ciliated cells of diverse organisms, as well as its connection to TRP channels, led us to hypothesize that it might play a role in left-right axis patterning. In *Xenopus*, left-right patterning requires leftward flow generated by motile cilia on the gastrocoel roof plate (GRP) to induce genes such as *Pitx2c* specifically in the left lateral plate mesoderm (Schweickert et al., 2010). In situ hybridization revealed that *Enkur* expression is restricted to the GRP in stage 17 embryos and ENKUR-GFP localized to the cilia of GRP cells (Figure 6A and B).

**Figure 6.**
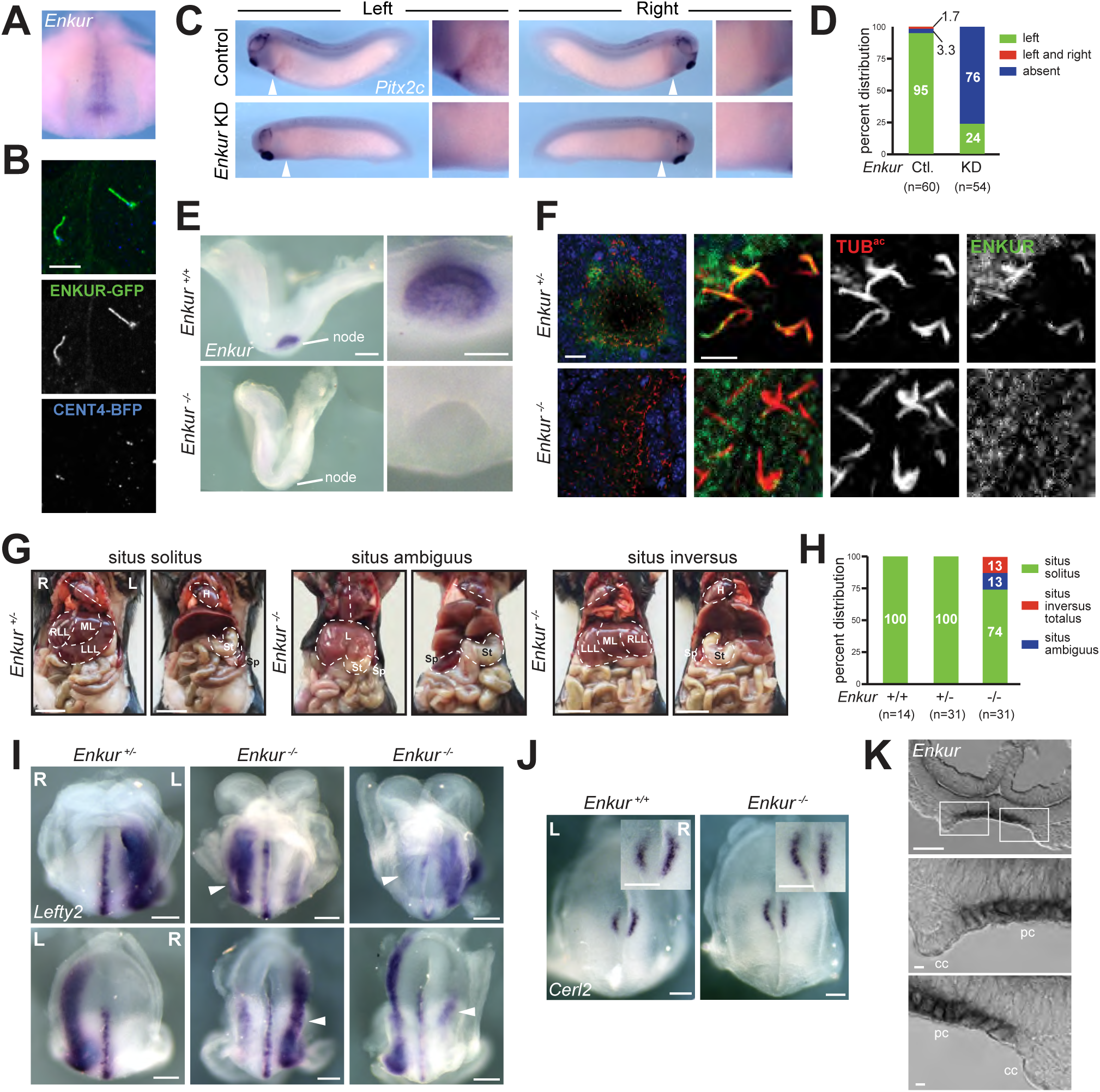
ENKUR is required for left-right axis patterning in mice. **(A)** Whole mount in situ hybridization of a stage 17 *X. laevis* embryo for *Enkur.* Ventral view of dorsal resection of embryo. Only the posterior region of the embryo is shown. *Enkur* is expressed in the gastrocoel roof plate (GRP). **(B)** Fluorescence imaging of *X. laevis* GRP cilia expressing ENKUR-GFP (green) and CENT4-BFP (blue). CENT4-BFP marks basal bodies. Scale bar, 5 μm. (**C**) In situ hybridization for *Pitx2c* of a stage 28 *X. laevis* control embryo and an *Enkur* knockdown (KD) embryo. *Pitx2c* is expressed in the left lateral plate (arrowhead and high magnification image) of control embryos but is absent in the *Enkur* KD embryo. **(D)** Quantification of *Pitx2c* expression patterns in control and *Enkur* KD embryos. **(E)** In situ hybridization of 2-4 somite stage littermate control and *Enkur*^-/-^ mouse embryos for *Enkur*. At this stage, *Enkur* is expressed exclusively in the node. *Enkur*^-/-^ embryos do not express detectable *Enkur*. Scale bar for left panel, 50μm. Scale bar for right panel, 25 μm. **(F)** Immunofluorescence imaging of 2-4 somite stage mouse embryos for ENKUR (green) and cilia (TUB^ac^, red). ENKUR localizes to the cilia of the node and is not detectable in the cilia of *Enkur*^-/-^ nodes. Nuclei are stained with Hoechst (blue). Scale bar for left panels, 10 μm. Scale bar for right 6 panels, 2.5 μm. **(G)** Photographs of thoracic and abdominal organ positions. The right (R) and left (L) side of the body are indicated. Some *Enkur*^-/-^ mice display situs ambiguus, illustrated here by abnormal heart apex orientation, midline liver and right-sided spleen. Some *Enkur*^-/-^ mice display situs inversus, a complete reversal of the left-right axis. Scale bar, 1 cm. **(H)** Quantification of situs in littermate control and *Enkur*^-/-^ mice. **(I)** In situ hybridization of 3-5 somite stage littermate control and *Enkur*^-/-^ embryos for *Lefty2*. Upper panels are rostral views. Lower panels are caudal views. The right (R) and left (L) side of the embryo are indicated. Control embryos exhibit *Lefty2* expression in the midline and left lateral plate mesoderm. *Enkur*^-/-^ embryos exhibit variable patterns of *Lefty2* expression, including expression predominantly in the right lateral plate mesoderm (middle) or partially in the right lateral plate mesoderm (right). Scale bar, 50 μm. **(J)** In situ hybridization of 3-5 somite stage littermate control and *Enkur*^-/-^ embryos for *Cerl2*. The right (R) and left (L) side of the embryo are indicated. Control embryos display an enrichment of *Cerl2* expression on the right side of the node. *Enkur*^-/-^ embryos express equal levels of *Cerl2* on both sides of the node. Scale bar, 100 μm. **(K)** Cryosectioned in situ hybridization of 2-4 somite stage mouse embryo for *Enkur*. Top panel depicts the entire node and lower two panels depict areas indicated by white boxes. *Enkur* is expressed predominantly within the pit cells (pc) and not the crown cells (cc) of the node. Scale bar top panel, 50 μm. Scale bar bottom two panels, 5 μm.

To determine if ENKUR function is required for establishing the left-right axis, we inhibited *Enkur* expression using a morpholino (Figure S5E). Whereas 95% of control embryos displayed *Pitx2c* in the left lateral plate mesoderm, *Pitx2c* expression was absent in 76% of *Enkur* morphants, suggesting that ENKUR is required for left-right axis patterning in *X. laevis* (Figure 6C and D).

The left-right axis defect in *Enkur* morphant *Xenopus* led us to investigate if the requirement for ENKUR in left-right patterning was conserved in mammals. We examined the developmental expression of mouse *Enkur* by situ hybridization, which revealed that, similar to *X. laevis*, 2-4 somite stage mouse embryos express *Enkur* exclusively in the node (Figure 6E and S5F). Immunofluorescent staining indicated that, again like *X. laevis,* ENKUR localizes to nodal cilia, suggesting that mouse ENKUR may function in mouse left-right patterning (Figure 6F).

We analyzed visceral organ placement in adult *Enkur*^-/-^ mice and found that 26% displayed abnormal situs (Figure 6G and H). Half of the affected mice had situs inversus totalis, a complete reversal of the left-right axis, and half displayed situs ambiguus, a variably abnormal positioning of organs that includes misorientation of the heart, right-sided stomach or spleen, asplenia, left-sided liver, and abnormal hepatic lobulation. Embryos that lack nodal cilia or have non-motile nodal cilia exhibit a higher incidence of abnormal situs than do *Enkur*^-/-^ embryos (Nonaka et al., 1998; Takeda et al., 1999). The presence of left-right axis defects in a minority of *Enkur*^-/-^ mice indicates that axis specification is not entirely randomized in the absence of ENKUR function.

An early event in left-right axis specification is the generation of leftward flow by nodal cilia which activates Nodal signaling specifically in the left lateral plate mesoderm (Collignon et al., 1996; Lowe et al., 1996; Saijoh et al., 2003). To determine if ENKUR participates in this step in left-right axis patterning, we examined the expression of *Lefty2*, a Nodal target gene (Meno et al., 1996). In situ hybridization revealed that in *Enkur*^-/-^ embryos, *Lefty2* expression in the lateral plate mesoderm was bilateral, and variably enriched on the left or right side (Figure 6I), indicating that ENKUR functions in an early step in left-right axis specification.

To test if ENKUR is acting upstream of the Nodal signaling pathway in left-right axis patterning, we analyzed the expression of *Cerberus like-2* (*Cerl2*), a secreted protein that antagonizes Nodal (Marques, 2004). In response to nodal fluid flow, *Cerl2* is downregulated on the left side of the node (Nakamura et al., 2012). In contrast to littermate control embryos, *Enkur*^-/-^ embryos displayed equal, bilateral expression of *Cerl2*, suggesting that ENKUR may be acting before Nodal signaling is activated, perhaps by generating or sensing leftward nodal flow (Figure 6J).

One model of nodal flow sensation contends that there are two types of monocilia present at the node: motile cilia of nodal pit cells generate flow and crown cell cilia on the periphery of the node detect the flow to distinguish left from right (Babu and Roy, 2013; McGrath et al., 2003). To investigate whether *Enkur* is expressed in one or both types of nodal cells, we examined *Enkur* expression in sectioned 2-4 somite stage embryos. *Enkur* was expressed by nodal pit cells but absent or only weakly expressed by crown cells (Figure 6K). The predominant expression of *Enkur* by pit cells suggests that ENKUR may contribute to leftward flow.

### Mutation of human *ENKUR* affects left-right axis specification

A hallmark of human ciliopathies affecting nodal cilia is situs inversus (Afzelius, 1976). We identified a consanguineous Azerbijaini kindred with two progeny that display situs inversus totalis (Figure 7A and B, S5G). Notably, neither affected individual exhibited recurrent airway infections or bronchiectasis, hallmarks of primary ciliary dyskinesia (PCD). Homozygosity mapping of the mother (OP-1605 I2) and one affected individual (OP-1605 II3) identified a short segment of homozygosity by descent on chromosome 10 containing *ENKUR* (Figure 7C). Whole exome sequencing of one affected sibling (OP-1605 II3) excluded previously identified mutations in all genes causative for PCD and heterotaxia but identified a homozygous variant, c.224-1delG, that alters the splice acceptor site of the second *ENKUR* intron (Figure 7D, Table S7). Sanger sequencing confirmed that this variant co-segregated with the phenotype and showed a recessive pattern of inheritance in the kindred (Figure 7E). Together, these data suggest that homozygous mutations in *ENKUR* are a cause of situs inversus in humans.

**Figure 7.**
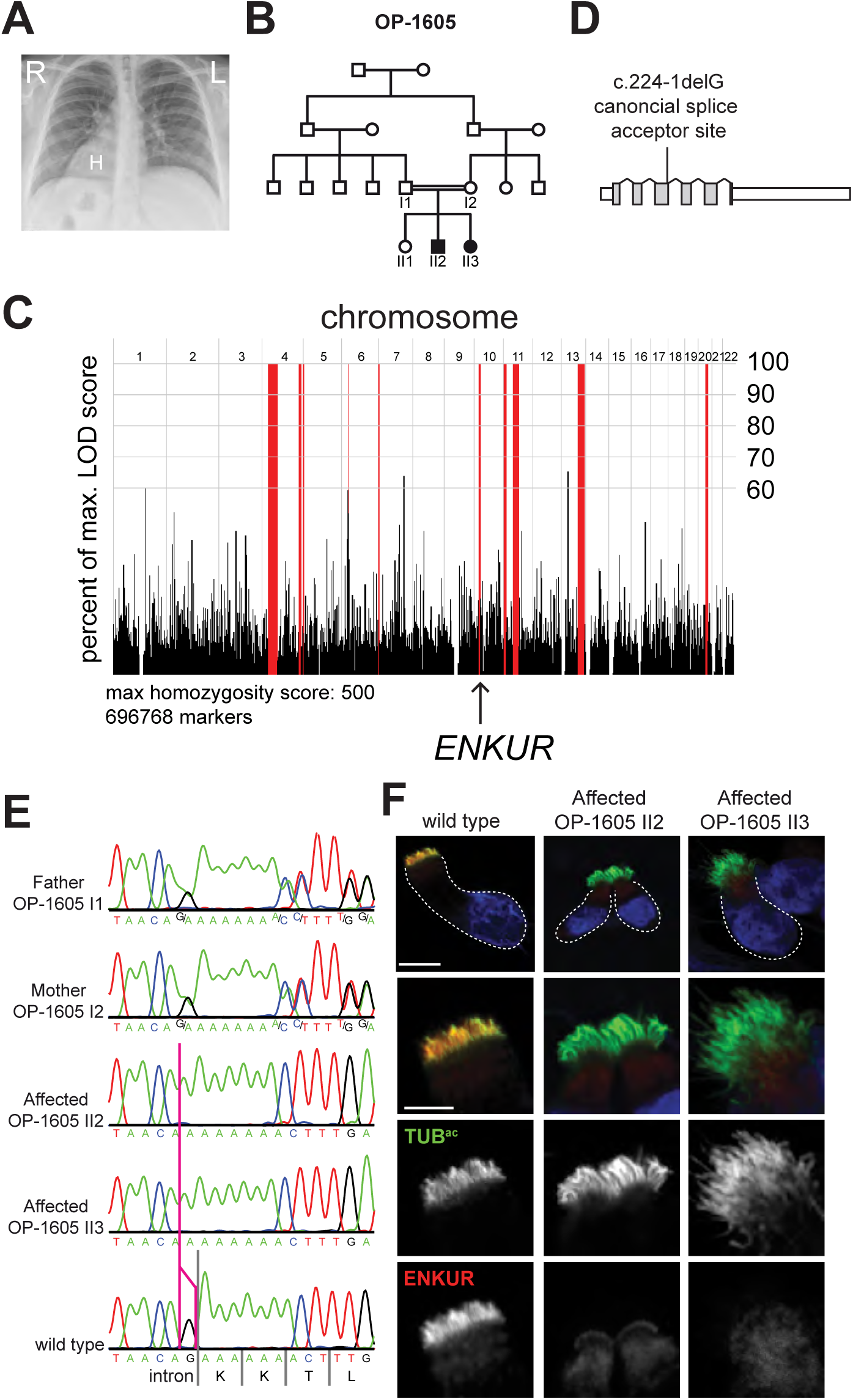
An inherited human *ENKUR* mutation causes situs inversus. **(A)** A chest X-ray of OP-1605 II3. The heart is indicated with an “H”, revealing dextrocardia. The right (R) and left (L) side of the body are indicated. (**B**) Pedigree of family OP-1605. Third degree consanguineous individuals are the parents of 2 siblings, individuals II2 and II3, affected with situs inversus. **(C)** Homozygosity mapping of the mother (OP-1605 I2) and one affected individual (OP 1605-II3) identified a short homozygous region on chromosome 10 containing *ENKUR*. **(D)** Schematic of the human *ENKUR* gene. The identified 1 bp deletion, c.224-1delG, occurs in the splice acceptor of the second intron. (**E)** Sanger sequence chromatograms of the *ENKUR* intron-exon boundary of the two affected individuals, their parents and an unrelated wild type individual. The pink line identifies the position of the last nucleotide of *ENKUR* intron 2 affected by the deletion. Both affected individuals are homozygous for c.2241-delG. Both parents are heterozygous for the mutation. **(F)** Immunofluorescence imaging of nasal epithelial cells from an unaffected control and two affected individuals for ENKUR (red) and cilia (TUB^ac^, green). Nuclei are stained with DAPI (blue). ENKUR localizes to cilia of control cells, and is missing from the cilia of affected individuals. The dotted line highlights the cell border. Scale bar for whole cells (top panel), 10 μm. Scale bar for cilia (bottom 3 panels), 5 μm.

To assess the effect of the homozygous mutation on ENKUR, we examined the motile cilia of nasal epithelial cells isolated from the respiratory tracts of both affected individuals and an unaffected control. We found that human ENKUR localizes to nasal epithelial cilia (Figure 7F). In contrast, neither individual with situs inversus exhibited ciliary localization of ENKUR (Figure 7F), suggesting that the human mutation abrogates ENKUR expression.

To determine whether ENKUR affects the localization of other ciliary proteins associated with situs inversus, we analyzed their localization in human nasal epithelia cells of an *ENKUR* mutant individual. Outer dynein arm (ODA) complex components DNAH5 and DNAH11, the ODA-docking complex component CCDC151, the inner dynein arm component DNALI1, the nexin-dynein regulatory complex component GAS8, the axonemal components CCDC11 and CCDC39, and the radial spoke component RSPH9 (Dougherty et al., 2016; Fliegauf et al., 2005; Hjeij et al., 2014; Kastury et al., 1997; LeDizet and Piperno, 1995; Merveille et al., 2011; Olbrich et al., 2015) all localized equivalently to the cilia of wild type control and homozygous *ENKUR* mutant cells (Figure S6). Likewise, ENKUR localized normally in human nasal epithelial cells from PCD-affected individuals with mutations in *CCDC40*, *CCDC11* or *DNAAF2* (Becker-Heck et al., 2011; Omran et al., 2008a) (Figure S7). Thus, ENKUR and several proteins associated with situs inversus are mutually dispensable for each other’s localization to cilia.

As neither homozygous *ENKUR* mutant individual exhibited the respiratory manifestations of PCD, we analyzed the motility of their nasal epithelial cell cilia. The cilia of both individuals with situs inversus displayed coordinated, wavelike beating, comparable to those of unaffected control individuals (Supplemental video S1 and S2). The ciliary beat frequencies of the affected individuals were 5.7±3.8 Hz and 5.6±1.3 Hz, within the normal range (Raidt et al., 2014) (Table S8). Whereas individuals with disrupted ciliary motility produce reduced levels of nasal nitric oxide (Lundberg et al., 1994), the nasal nitric oxide production rates of the two individuals with situs inversus were 155 ml/min and 234 ml/min, exceeding the 77 ml/min production rate below which is consistent with PCD (Table S8). According to these measures, the respiratory ciliary function of both individuals with homozygous *ENKUR* mutations was normal.

To determine if, like humans, ENKUR is dispensable for ciliary motility in the respiratory tract of mice, we analyzed airway cilia of *Enkur*^-/-^ mice. We analyzed the beating of cilia on tracheal epithelial cells of *Enkur*^-/-^ mice and discovered that, like humans, the motility of mutant cilia was not distinguishable from wild type (Supplemental video S3 and S4).

Together, these data demonstrate that ENKUR is a highly conserved ciliary protein that is not required for ciliary motility in the airway, but that is required for left-right axis determination in *Xenopus* and mice. Furthermore, when mutated, *Enkur* may be a cause of situs inverus in humans.

## DISCUSSION

Communication of environmental or intercellular information is a critical function of the cilia of organisms from diverse phyla throughout Eukarya. In vertebrates, the signaling functions of cilia are especially important for sight, olfaction and development. At what points in animal evolution did different signaling pathways become associated with cilia? Whereas the structural proteins that make up cilia are conserved among most ciliated Eukaryotes, including humans, few metazoan signaling proteins are conserved outside of the animal kingdom. Therefore, we identified ciliary proteins from organisms representing major steps in animal evolution: choanoflagellates, whose phylogenetic position informs the origin of multicellular animals, sea anemones, radially symmetric sister group to bilaterally symmetric animals, and sea urchins, basal Deuterostomes that provide insights into the ancestor of chordates.

The ciliomes indicate that proteins involved in GPCR and Hh signaling were associated with cilia before bilateria arose and TRP channels were associated with cilia before the emergence of animals. Many of the identified proteins with homologs in mammals, the conserved ciliome, had a phylogenetic distribution paralleling the presence of cilia: present in ciliated organism of the Animal, Plant, Fungi and Protist kingdoms but lost from unciliated organisms within these groups. These ciliary proteins are therefore likely to have been present in the last common ancestor of all extant eukaryotes, a ciliated organism.

One protein that we identified in the ciliomes of choanoflagellates, sea anenomes and sea urchins, suggesting that it became associated with ciliary function before the emergence of animals, was ENKUR, a highly conserved protein found in diverse ciliated eukaryotes. We found that *Enkur* is expressed in tissues with motile ciliated cells, including those with single cilia (*i.e., Xenopus* and mouse nodal cells, sea anemone and sea urchin embryonic epithelial cells) and those with multiple cilia (*i.e.*, mouse tracheal epithelial cells, *Xenopus* embryonic epidermal cells). In addition, ENKUR from animals and choanoflagellates localizes to cilia, raising the possibility that it acts within cilia to generate or respond to fluid flow. Mutation of *Enkur* in the mouse did not compromise ciliary beat frequency in tracheal epithelial cells, indicating that, unlike many ciliary proteins with expression patterns and phylogenetic distributions similar to ENKUR (*e.g.*, outer arm dyneins), it is not essential for ciliary motility. Instead, ENKUR is required for patterning of the left-right axis in both *Xenopus* and mouse, suggesting that it is required for some aspect of ciliary beating or signaling in the embryonic node. Significantly, humans with a homozygous mutation in *ENKUR* displayed inversion of their left-right axis, but no signs of defective mucociliary clearance, suggesting that human *ENKUR* loss of function does not cause PCD, but may be a cause situs inversus.

ENKUR has previously been shown to interact with TRP channels and calcium signaling proteins, suggesting that it may bridge the gap between TRP channels and downstream signaling components (Sutton et al., 2004). Calcium signaling and the ciliary TRP channel PKD2 have been implicated in left-right axis patterning, although their exact roles remain unclear (McGrath et al., 2003; Pennekamp et al., 2002; Yoshiba et al., 2012). In the sea urchin, PKD2 is expressed at the tip of the archenteron (Tisler et al., 2016), where we find *Enkur* to also be expressed. The archenteron tip possesses motile cilia that beat rotationally, similar to cilia of the node, although how cilia in sea urchin left-right axis patterning function remains a subject of investigation (Tisler et al., 2016; Warner et al., 2016). PKD2 has also been shown to be expressed in the ciliary tuft of sea urchin embryos, a structure hypothesized to have sensory function (Tisler et al., 2016). We found that, like PKD2, *Enkur* mRNA is enriched in the cells of both the sea urchin and sea anemone ciliary tuft. It will be of great interest to assess whether ENKUR regulates or interprets calcium responses in nodal, archenteron or tuft cilia.

In addition to ENKUR, we identified 130 other proteins in the ciliomes of all three organisms examined here. As expected, these proteins included many recognized to have evolutionarily ancient ciliary roles, including structural components (*e.g.,* Tektin-1, β-Tubulin), intraflagellar transport (*e.g.,* IFT57, IFT80, IFT81, IFT122, IFT172) and ciliary movement (*e.g.,* axonemal dyneins, Nexin-Dynein regulatory complex components, radial spoke proteins, SPAG17, HYDIN). Other highly conserved ciliary proteins, such as components of the transition zone and basal body, were not detected as our methods amputated cilia distal to the basal bodies and transition zones (Sanders and Salisbury, 1994; Stephens, 1995). Similarly, we did not detect ciliopathy-associated proteins that are not ciliary components, such as those PCD-associated proteins that function within the cytoplasm to assembly dynein arms (Kott et al., 2012; Loges et al., 2009; Mitchison et al., 2012; Omran et al., 2008b). Like ENKUR, we predict that many of the uncharacterized or poorly characterized proteins present in the ciliomes of all three organisms (*e.g*., C9orf116, C4orf22, C10orf107, C5orf49) will localize to cilia and have ciliary functions.

The sea anemone and sea urchin ciliomes share 250 proteins not identified in the choanoflagellate ciliome. Some of these proteins, such as EVC2, have functions associated with Hh signaling, a pathway present in many multicellular animals including sea anemones and sea urchins but absent, at least in its canonical form, from choanoflagellates (Adamska et al., 2007). EVC2 is part of a complex comprised of four proteins, EVC, EVC2, IQCE and EFCAB7, each of which were detected in at least one ciliome (Pusapati et al., 2014). Orthologs of the genes encoding these proteins are not present in the choanoflagellate genome, suggesting that they emerged during animal evolution in parallel with the development of ciliary Hh signaling.

In addition to EVC and EVC2, we identified a structurally related protein called EVC3 in the ciliomes of sea urchins, sea anemone and choanoflagellates. The phylogenetic distribution of the EVC complex components and EVC3 suggests that EVC3 may represent an ancestral ciliary protein, with EVC and EVC2 emerging in multicellular animals as regulators of Hh signaling (Pusapati et al., 2014). The identification of EVC3 proteins in cilia suggests that EVC proteins likely have ancient functions at cilia where EVC3 presumably had Hh-independent functions.

Other known ciliary proteins such as TULP3, a regulator of Hh signaling, were identified exclusively in the sea urchin ciliome. Although homologs of TULPs are present in the genomes of both choanoflagellates and sea anemones, an ortholog of TULP3 is missing, suggesting that the TULP family expanded in animals and TULP3 may be a Deuterostome innovation.

Beyond Hh signaling components, we detected proteins involved in other pathways, such as GPCR signaling, in the ciliomes of both sea anemones and sea urchins. Homologs of four mouse GPCRs were present in the sea urchin or sea anemone ciliome. In addition to GPCRs, orthologs of multiple G protein components were identified in these ciliomes, including GNAS, GNAQ, GNAO1 and GNB2. The roles of the G protein Transducin in phototransduction at the photoreceptor outer segment (a modified cilium) and of the G protein GNAL in olfaction at the cilia of olfactory sensory neurons are well appreciated (Gießl et al., 2004; Kuhlmann et al., 2014). The identification of multiple G proteins in the ciliomes of sea anemones and sea urchins suggests that their association with cilia arose early in animal evolution. The co-existence of GPCR and Hh signaling proteins in cilia of early-branching animals may have facilitated the evolution of crosstalk between the two pathways, exemplified by the involvement of the GPCR GPR161, β-Arrestins and GRKs in mammalian Hh signal transduction (Chen et al., 2011; Evron et al., 2011; Mukhopadhyay et al., 2013; Pal et al., 2016). As the ciliomes contained previously identified members of Hh, GPCR and TRP channel signaling, we hypothesize that they also contained unrecognized ciliary regulators of these signaling pathways.

The ciliomes identified 73 proteins encoded by genes that, when mutated, cause human ciliopathies such as PCD (*e.g., HYDIN, CCDC39, TTC25*), Bardet-Biedl syndrome (*e.g., BBS1*, *BBS2*, *BBS3/ARL6*), retinal degeneration (*e.g., LCA5, ARL3, CLUAP1)* and nephronophthisis (*e.g.*, *DCDC2A, TRAF3IP1*) (Hildebrandt et al., 2011). Another 241 genes represented in these ciliomes are associated with human diseases with no established connection to cilia. Perhaps some of these diseases are caused by defects in cilia, as discussed for WDR65 and van der Woude syndrome, above. Beyond WDR65, we identified RAB28 in the sea anemone ciliome, mutations in which are associated with a form of retinitis pigmentosa called progressive rod-cone dystrophy 18 (Cord18) (Roosing et al., 2013). Therefore, we hypothesize that Cord18 is caused by RAB28-associated dysfunction of the photoreceptor outer segment. Similarly, we identified NEK9, in the ciliome, which is linked to a form of skeletal dysplasia recently proposed to be a ciliopathy (Casey et al., 2016). We predict that, like ENKUR and situs inversus, other ciliome members identified here will prove to underlie orphan ciliopathies.

The identification of an extensive compilation of ciliary proteins from organisms at key phylogenetic nodes has shed light on the evolutionary history of ciliary signaling proteins, suggesting that ciliary signaling pathways, such as Hh and GPCR signaling, evolved before the emergence of Bilaterians or, in the case of TRP channel signaling, before the emergence of animals. Using evolutionary proteomics, we identified previously uncharacterized ciliary proteins, including ENKUR, which our data suggest may underlie a human disease, demonstrating promise for further studies of ciliome members and for future applications of evolutionary proteomics. The use of the ciliomes to identify an evolutionarily conserved function for ENKUR illustrates how a protein’s evolutionary association with an organelle can help elucidate its function.

Our cilia proteomics approach has provided insights into the evolutionary history of ciliary signaling proteins and identified previously unknown cilia proteins. Similar to the analysis brought to bear on cilia here, we propose that proteomic profiling of other organelles, such as mitochondria or lysosomes, from multiple, diverse organisms will help to define their evolutionary trajectories and elucidate their core components.

## METHODS

### Animal husbandry and embryo culture

#### Sea urchins

Adult *S. purpuratus* were obtained from UC Davis Bodega Marine Laboratory (Bodega Bay, CA) or Kerckhoff Marine Laboratory (Corona Del Mar, CA) and housed in tanks containing artificial seawater (Instant Ocean, Blacksburg, VA) at 12-15°C. Gametes were collected by bathing the gonads with 0.5M KCl to stimulate release of eggs and sperm. Eggs were washed 3 times with natural seawater and approximately 2,000,000 eggs were fertilized with a drop of sperm in 500ml of natural seawater. The embryos were cultured at approximately 14°C with constant, gentle stirring.

#### Sea anemones

*N. vectensis* were maintained in 33% artificial seawater and spawning was induced as described (Fritzenwanker and Technau, 2002). Embryos were cultured at 21°C in 33% artificial seawater.

#### Choanoflagellates

An environmental isolate of *S. rosetta* (ATCC50818) that had been treated with antibiotics to reduce bacterial diversity and subsequently supplemented with *Algoriphagus machipongonensis* to induce colonies (Fairclough et al., 2010) was cultured in cereal grass infused artificial seawater (Tropic Marine, Montague, MA) diluted to 10% in artificial seawater (King et al., 2009) at approximately 22°C.

#### Zebrafish

*Danio rerio* of the *Ekkwill* strain were reared at 28°C and embryos were obtained from natural matings. Embryos were cultured at 28°C for 4-6 hours post fertilization and then maintained at approximately 22°C.

#### Xenopus laevis

*X. laevis* husbandry and experiments were performed using standard conditions and following animal ethics guidelines of the University of Texas at Austin, protocol number AUP-2015-00160.

#### Nematostella vectensis

Animals were maintained in 1/3 filtered seawater and induced to spawn as described (Fritzenwanker and Technau, 2002). Egg packages were fertilized and embryos were raised at 21°C.

### Cilia isolation

#### Sea urchins

Gastrula stage *S. purpuratus* embryos were concentrated by centrifugation at 170-200 g for 4-10 minutes at 4°C and washed 3-4 times with artificial seawater. To amputate cilia, embryos were gently resuspended in a high salt solution (artificial seawater + 0.5M NaCl) approximately 10 times the volume of the embryo pellet. The samples were immediately spun at 400 g for 5 minutes at 4°C to pellet deciliated embryos. The supernatant was transferred to a fresh tube and spun a second time to pellet any remaining embryos. The supernatant containing cilia was then spun at 10,000 g for 20 minutes at 4°C and the resulting cilia pellet was collected for subsequent analysis. Cilia samples for mass spectrometry were resuspended in 0.1% *Rapi*Gest SF Surfactant (Waters, Milford, MA).

To separate the axonemes from the membrane plus matrix fraction, pelleted cilia were resuspended in extraction buffer (1% Nonidet NP-40, 30mM Hepes, pH7.4, 5mM MgSO4, 0.5mM EDTA, 25mM KCl, 1mM DTT) and centrifuged at 30,000 g for 20 minutes at 4°C. The supernatant (membrane plus matrix fraction) was separated from the pellet (axonemes) and the pellet was resuspended in SDS extraction buffer (0.5% SDS, 50mM NaCl, 50mM Tris, pH 7.4). Both samples were precipitated in methanol: chloroform (4:1) and resuspended in 0.1% *Rapi*Gest SF Surfactant for mass spectrometry.

#### Sea anemones

*N. vectensis* planula larva (day 3 of development) were washed twice with 33% artificial seawater containing 20mM DTT and then transferred to high salt solution (artificial seawater, 0.5M NaCl, 100mM DTT, Complete Mini protease inhibitor without EDTA (Roche)) and incubated with gentle agitation at 4°C for 5 minutes. The embryos were removed by centrifugation at 400 g for 2 minutes at 4°C. The supernatant containing cilia was transferred to LoBind microcentifuge tubes (Eppendorf, Hamburg, Germany) and centrifuged at 10,000 g for 10 minutes at 4°C to pellet cilia. The pellet was gently resuspended and diluted to 600μl with 33% artificial seawater containing 100mM DTT. The cilia sample was overlain onto a sucrose step gradient from 80-30% sucrose (sucrose solution contained 50mM Tris, pH8.0, 10mM DTT), step size 10%, 1ml per step. The sample was spun at 100,000 g in a swinging bucket rotor for 3 hours at 4°C and fractions were collected from the bottom of the tube by puncturing with a 30 gauge needle and allowing the sample to flow by gravity. A portion of each fraction was analyzed by immunoblot to determine which fractions contained high levels of TUB^ac^ and low levels of Actin (Figure 1C, e.g. fractions 6 and 7). To concentrate the samples and remove the sucrose, the protein was precipitated using 16% trichloroacetic acid (TCA) and washed 2 times with acetone, then resuspended in *Rapi*Gest SF Surfactant for mass spectrometry.

#### Choanoflagellates

Before flagellar amputation, 8L of *S. rosetta* culture were harvested at peak density by centrifugation at 4,000 g for 15 minutes at 4°C to pellet colonies. Colonies were concentrated into approximately 250ml and passed through a 40 μm filter to remove clumps of bacteria. *S. rosetta* were then centrifuged at 2,000 g for 15 minutes at 4°C. The pellet was resuspended in 20ml of artificial seawater (Tropic Marin). The sample was split in half and overlain onto 2ml of Percoll solution (8% Percoll - GE Healthcare, 0.5M sorbitol, 50mM Tris, pH8.0, 15mM MgCl_2_, 1% artificial seawater) and spun at 1,000 g for 10 minutes at room temperature. The supernatant containing bacteria was removed and the pellet of *S. rosetta* colonies was gently resuspended in 50ml of artificial seawater. The culture was incubated for 20 hours to allow the *S. rosetta* to eat the residual bacteria.

To amputate flagella, *S. rosetta* colonies were concentrated by centrifugation at 2,000 g for 15 minutes and resuspended in 10 ml of HMSA (10mM Hepes, pH7.4, 5mM MgSO_4_, 10% sucrose, 5% artificial seawater). The colonies were transferred to a 10cm dish and 700μM dibucaine was added. The dish was agitated gently for 5 minutes and deflagellation was visually followed using a stereomicroscope. Additional dibucaine was added as needed until the majority of flagella were amputated. Immediately after flagellar amputation, 5ml of 10mM Hepes, pH7.4, 5mM MgSO_4_, 1mM EGTA was added to the sample.

To separate the flagella from the cell bodies, the sample was split in half and purified over Percoll as described above. The top, clear layer containing the flagella was collected and the pellet and bottom 2 ml containing most of the cells were discarded. The flagellar fraction was spun at 16,000 g for 20 minutes at 4°C to pellet the flagella. To further purify the sample, the flagellar pellet was gently resuspended in ~800μl of artificial seawater and overlain onto a sucrose step gradient. The sucrose was dissolved in 10mM Hepes, pH7.4, 5mM MgSO_4_, 5% artificial seawater and a step gradient from 40-80% sucrose was prepared using 10% steps, 1ml of each step. The sample was spun at 100,000 g for 3 hours at 4°C. To collect fractions, a hole was punctured in the bottom of the tube using a 25 gauge needle and fractions between 100 and 700μl were captured.

A small portion of each fraction was analyzed by immunoblot to determine which fractions contained high levels of TUB^ac^ and low levels of Actin (Figure 1G, Fractions 4 and 5). Samples from 2 experiments were pooled for mass spectrometry. Fraction 1 and 10 were also analyzed by mass spectrometry to use for comparison (see below). The samples were TCA precipitated, washed and resuspended for mass spectrometry as described above.

### Mass spectrometry and protein identification

Purified cilia or flagella were lysed in a buffer containing 8M urea, 150mM NaCl, and protease inhibitors (Complete, EDTA-free tablet, Roche) and digested with Trypsin (Ramage et al., 2015). For samples that were fractionated, the digested peptides (100μg) were fractionated using hydrophilic interaction chromatography (HILIC). The samples were injected onto a TSKgel amide-80 column (Tosoh Biosciences, 2.0 mm x 15 cm packed with 5 μm particles) equilibrated with 10% HILIC buffer A (2% acetonitrile [ACN], 0.1% trifluoroacetic acid [TFA]) and 90% HILIC buffer B (98% ACN, 0.1% TFA) using an AKTA P10 purifier system. The samples were then separated using a one-hour gradient from 90% HILIC buffer B to 55% HILIC buffer B at a flow rate of 0.3 ml/min. Fractions were collected every 1.5 min and combined into 12 fractions based on the 280 nm absorbance chromatogram. Fractions were evaporated to dryness and reconstituted in 20 μl of 0.1% formic acid for mass spectrometry analysis.

All samples were analyzed on an Orbitrap Elite mass spectrometry system equipped with an Easy-nLC 1000 HPLC and autosampler. HILIC fractionated samples were injected onto a pre-column (2 cm x 100 μm I.D. packed with 5 μm C18 particles) in 100% buffer A (0.1% formic acid in water) and separated on an analytical column (10 cm x 75 μm I.D. packed with 1.9 μm C18 particles) with a 60 minute reverse phase gradient from 5% to 30% buffer B (0.1% formic acid in 100% ACN) at a flow rate of 400 nl/min. The mass spectrometer continuously collected spectra in a data-dependent manner, acquiring a full scan in the Orbitrap (at 120,000 resolution with an automatic gain control target of 1,000,000 and a maximum injection time of 100 ms), followed by collision-induced dissociation spectra for the 20 most abundant ions in the ion trap (with an automatic gain control target of 10,000, a maximum injection time of 10 ms, a normalized collision energy of 35.0, activation Q of 0.250, isolation width of 2.0 m/z, and an activation time of 10.0). Singly and unassigned charge states were rejected for data-dependent selection. Dynamic exclusion was enabled to data-dependent selection of ions with a repeat count of 1, a repeat duration of 20.0 s, an exclusion duration of 20.0 s, an exclusion list size of 500, and exclusion mass width of +/- 10.00 parts per million (ppm).

Samples that were not HILIC fractionated were injected onto a high resolution C18 column (25 cm x 75 μm I.D. packed with ReproSil Pur C18 AQ 1.9 μm particles) in 0.1% formic acid and then separated with a two-hour gradient from 5% to 30% ACN in 0.1% formic acid at a flow rate of 300 nl/min. The mass spectrometer collected data in a data-dependent fashion, collecting one full scan in the Orbitrap at 120,000 resolution with an AGC target of 1,000,000 followed by 20 collision-induced dissociation MS/MS scans in the dual linear ion trap with an AGC target of 30,000 for the 20 most intense peaks from the full scan. Dynamic exclusion was enabled to exclude masses within +/- 10 ppm for 30 s a repeat count of 1. Charge state screening was employed to reject analysis of singly charged species or species for which a charge could not be assigned.

Raw mass spectrometry data were analyzed by the Protein Prospector suite (Clauser et al., 1999). Data were matched to *S. purpuratus*, *N. vectensis* and *S. rosetta* protein sequences downloaded from UniProt containing 28,593, 24,435, and 11,698 protein sequences, respectively, concatenated to a decoy database where each sequence was randomized in order to estimate the false positive rate. The searches considered a precursor mass tolerance of +/- 20 ppm and fragment ion tolerances of 0.8 da, and considered variable modifications for protein N-terminal acetylation, protein N-terminal acetylation and oxidation, glutamine to pyroglutamate conversion for peptide N-terminal glutamine residues, protein N-terminal methionine loss, protein N-terminal acetylation and methionine loss, and methionine oxidation, and constant modifications for carbamidomethyl cysteine. Prospector data was filtered using a maximum protein expectation value of 0.01 and a maximum peptide expectation value of 0.05.

### Membrane plus matrix enrichment analysis

To identify proteins that were enriched in the membrane + matrix fraction of sea urchin cilia, the unique peptide count (the number of distinct peptide sequences identified for each protein) was used as a proxy for protein abundance. [M/M+1/total+1]-[Ax+1/total+1] = enrichment score. The membrane plus matrix enrichment score for cilia from early gastrula embryos and late gastrula embryos was summed to determine the final enrichment score.

### Choanoflagellate ciliome subtractive analysis

To identify proteins that were enriched in the cilia of *S. rosetta,* we compared the unique peptide count of two cilia samples (Figure 1G, fractions 4 and 5) to the cell body sample (fraction 1) and the microvilli sample (fraction 10) and discounted proteins that did not have a unique peptide count in either fraction 4 or 5 that was greater than fraction 1 and 10.

### Identification of the conserved ciliome

To identify ciliome proteins with a homolog in *M. musculus*, we used the BLAST+ suite of executables (Camacho et al., 2009). For each organism analyzed by mass specrometry, full predicted protein sequences of proteins associated with 2 or more unique peptide hits were used to query the *M. musculus* proteome (ensembl.org) using blastp (Expected value cutoff 1e^-5^). The top hit was classified as the mouse homolog. If reciprocal blast of the mouse protein homolog to the source organism’s proteome (downloaded from unprot.org) retrieved the original query protein (best reciprocal BLAST method), we considered the two proteins to be orthologs.

### CRAPome analysis

To remove common protein contaminants from the conserved ciliome, we used the CRAPome database (crapome.org) (Mellacheruvu et al., 2013) to determine the number of times each protein was identified in another control proteomics experiment. We removed proteins from the conserved ciliome that were identified in over 70 control experiments (Table S1).

### SYSCILIA gold standard analysis

To determine the proportion of ciliary SYSCILIA gold standard proteins (SCGSv1) (van Dam et al., 2013), with localization distal to the transition zone, that was present in the ciliome of each organisms, we determined the number of human ciliary SCGSv1 proteins had an ortholog in the proteome of *S. purpuratus*, *N. vectensis* and *S. rosetta* using InParanoid8 (http://inparanoid.sbc.su.se/cgi-bin/index.cgi) (Sonnhammer and Östlund, 2015). We then counted the number of proteins in the ciliome of each organism with an ortholog to a human ciliary SCGSv1 protein. To calculate the percentage of ciliary SCGSv1 proteins in previously published ciliomes, we downloaded data sets from Cildb V2.0 using the low stringency threshold (Avidor-Reiss et al., 2004; Ishikawa et al., 2012; Li et al., 2004, Mick et al., 2015; Ostrowski et al., 2002; Pazour et al., 2005).

### Choanoflagellate-*Chtamydomonas* ciliome comparison

We used blastp (Expected value cutoff 1e^-5^) to determine which protein homologs in the *S. rosetta* ciliome were present in the *Chlamydomonas reinhardtii* ciliary proteome (Pazour et al., 2005). Proteins with 2 or more peptides in the *C. reinhardtii* ciliary data set were included in the analysis.

### Identification of known ciliary proteins and disease classification

To identify the ciliome proteins previously implicated to be involved in ciliary biology, the Biomart tool in Cildb V2.0 (cildb.cgm.cnrs-gif.fr) was used to obtain the mouse ortholog of all cilia in 15 studies using the medium threshold (Avidor-Reiss et al., 2004; Blacque et al., 2005; Boesger et al., 2009; Broadhead et al., 2006; Cao et al., 2006; Chen et al., 2006; Efimenko et al., 2005; Laurençon et al., 2007; Li et al., 2004; Liu et al., 2007; Mayer et al., 2008; Merchant et al., 2007; Ostrowski et al., 2002; Pazour et al., 2005; Smith et al., 2005). For 4 studies that were not included in Cildb but used in our comparison, we obtained protein identifiers directly (Choksi et al., 2014; Ishikawa et al., 2012; Mick et al., 2015; Narita et al., 2010). See Table S4 for details on each study.

Online Mendelian Inheritance in Man (OMIM; omim.org), Orphanet (orpha.net) and Developmental Disorders Genotype-to-Phenotype database (DDG2P) (Wright et al., 2015) databases were used to identify genes associated with human disease.

### CLIME analysis

The phylogenetic distributions of ciliome members with mouse homologs was assessed by a computational algorithm, CLustering by Inferred Models of Evolution (CLIME) (Li et al., 2014). We evaluated how closely the ciliome member matched the phylogenetic distribution of cilia by assigning it a positive point for every ciliated organism in which it has a detected homolog (listed in Figure S2 legend), and a negative point for every unciliated organism in which it has a detected homolog, with a prokaryote homolog being weighted tenfold more. Genes that scored greater than 29 in this metric were considered to be co-evolved with cilia. Genes that scored less than 1 were considered to be not co-evolved with cilia. Genes with intermediate scores were considered to be neither.

### Protein sequences alignments

Alignments were built using MUSCLE and displayed using Jalview. The ClustalX color scheme was used to indicate residues with similar chemical properties.

### Random selection of proteins for ciliary localization screen

To select proteins for subcellular localization studies, 49 proteins from the high-confidence ciliome (Table S1) and 30 human genes not present in the ciliomes were selected by the RANDBTWN function of Excel (Microsoft) from the ENSEMBL list of human protein coding genes. Of these genes, 40 were represented by Gateway-compatible clones of the human ortholog present in the hORFeome v8.1 (Yang et al., 2011). We used Gateway-mediated subcloning to insert the open reading frame of the remaining genes in frame with the EGFP open reading frame of pCS-EGFP-DEST (Villefranc et al., 2007), as described below in Expression constructs and *in situ* probe plasmid generation. The list of the genes cloned is included in Table S5.

### Mammalian cell line culture and transfection

RPE-1 cells (ATCC CRL-4000) were cultured in Dulbecco’s modified Eagle’s medium (DMEM) High Glucose/F-12 50:50 (Cell Culture Facility (CCF)), University of California, San Francisco, CA) supplemented with 10% fetal bovine serum (FBS) (Gibco) and Glutamax (Gibco). IMCD3 cells (ATCC CRL-2123) were cultured in DMEM:F12 (Gibco) supplemented with 10% FBS and Glutamax. All cell lines were transfected using jetPRIME (Polyplus-stransfection, New York, NY).

### Mouse tracheal epithelial cell culture and infection

Mouse tracheal epithelial cell (mTECs) were isolated from C57BL/6J mice and cultured as described previously (You and Brody, 2012). Proliferating mTECs were infected with lentivirus carrying shRNAs (produced by UCSF Viracore, San Francisco, CA) as described previously (Vladar and Brody, 2013). To select for infected cells, mTECs were cultured with 2- 4mg/ml Puromycin for 10 days before initiating differentiation by culturing cells at the air-liquid interface. See Table S9 for shRNA sequences.

### Immunofluorescent staining and imaging

#### Sea urchin embryos

*S. purpuratus* embryos were fixed in 2% paraformaldehyde (PFA) in PEM (20mM PIPES, 0.5mM EGTA, 20mM MgCl_2_) + 0.1% Triton X-100 for 3 days, blocked in PBST (PBS + 0.1% Triton X-100) containing 10% donkey serum for 1 hour, and incubated with primary antibodies overnight at 4°C. Embryos were incubated with secondary antibodies and Hoechst 33342 (1μg/ml) for 1 hour, then incubated for 10 minutes with increasing concentrations of glycerol (10%, 25%, 50% in PBS; 10 minutes for each glycerol solution). Embryos were mounted in 50% glycerol in PBS for imaging.

#### Sea anemone embryos

To visualize Actin and TUB^ac^, *N. vectensis* embryos were fixed for 1.5 minutes in 3.7% Formaldehyde, 0.25% Gluteraldehyde in 33% artificial seawater and then in 3.7% Formaldehyde, 0.1% Tween-20 for 1 hour at 4°C. Embryos were blocked in PBST containing 10% donkey serum for 1 hour and subsequently incubated with primary antibodies and Alexa Fluor 555 Phalloidin (Invitrogen) overnight at 4°C. After washing with PBST, the embryos were incubated with secondary antibodies and DAPI (1μg/ml) for 1 hour. Primary and secondary antibodies were diluted in PBST + 2% BSA. Embryos were mounted in ProLong Gold Antifade Mounant (Invitrogen).

#### Choanoflagellate colonies

*S. rosetta* colonies were Percoll purified as described above to remove bacteria, placed on poly-L-lysine coated coverslips and incubated for 15 minutes to allow cells to attach. 6% acetone diluted in PEM was applied to the cells for 5 minutes, removed and the cells were subsequently fixed in 4% PFA in PEM for 10 minutes. Cells were blocked with 2% BSA in PEM + 0.1% Triton X-100. To stain for Actin and β-tubulin, embryos were incubated with primary antibodies overnight and secondary antibodies, Alexa Fluor 568 Phalloidin (Invitrogen) and DAPI (1μg/ml) for 1 hour. Coverslips were mounted in ProLong Gold Antifade Mounant (Invitrogen).

#### Mammalian cells

Cells were cultured on coverslips, fixed for 8-10 minutes with 4% PFA in PBS, and blocked with 5% Donkey Serum in PBST for 30 minutes. Cells were incubated with primary antibodies overnight at 4°C and secondary antibodies and Hoechst 33342 for 1 hour. All antibodies were diluted in 2% BSA, 1% donkey serum in PBST. Coverslips were mounted in ProLong Gold or Diamond Antifade mountant (Invitrogen).

#### Isolated cilia

Isolated cilia were applied to poly-L-lysine coated coverslips and incubated for 15 minutes to allow cilia to attach to the surface of the coverslip, then fixed with 4% PFA in PBS for 10 minutes and immunostained as described for mammalian cells.

#### Mouse embryos

Pregnant female mice were sacrificed, embryos isolated in PBS and fixed in 4% PFA in PBS for 30 minutes. The embryos were washed with PBST three times, 5 minutes each. After blocking in 1% BSA and PBST for two hours, the embryos were incubated in primary antibodies overnight in blocking solution at 4°C. Following washes with PBST, the samples were incubated with secondary antibodies and Hoechst 33342 in blocking solution for 1 hour. The embryos washed with PBST and mounted in ProLong Diamond Antifade mountant (Invitrogen).

#### mTECs

mTEC cells were fixed on culture membranes in 4% PFA in PBS for 10 minutes. The cells were incubated in 1% SDS, PBST for 5 minutes, washed with PBST and then stained as described above for mammalian cells. Structured illumination microscopy (SIM) data for mTECs (Figure 5H) were collected on a Nikon N-SIM microscope in 3D-SIM mode using an Apo TIRF 100x/1.49 Oil objective.

#### Zebrafish

*D. rerio* embryos were fixed in 4% PFA in PBS for 2 hours, washed 3 times in PBS, blocked in PBDT (PBS + 1% BSA, 1% DMSO, 0.5% Triton X-100) containing 10% donkey for 1 hour serum and stained with primary antibody diluted in PBDT overnight at 4°C. Embryos were incubated with secondary antibodies and Hoechst 33342 (1μg/ml) for 1 hour and mounted in 50% glycerol for imaging.

#### Human nasal brush biopsies

Nasal brush biopsies were obtained from the middle turbinate with a cytology brush and suspended in RPMI cell culture medium (GIBCO), as previously described (Wallmeier et al., 2014). Cells were spread on glass slides, air dried and stored at -80°C until use. Defrosted samples were fixed in 4% PFA in PBS for 15 minutes. After washing with PBS, the cells were permeabilized with 0.2%Triton-X 100 in PBS for 15 minutes. Following washes with PBS, samples were blocked in blocking solution (1-5% skim milk in PBS) at 4°C overnight. Samples were incubated for at least 3 hours with primary antibodies diluted in blocking solution. Slides were rinsed with PBST, washed with blocking solution and incubated with secondary antibodies in blocking solution for 30 minutes. After washes with PBS, the nuclei were stained with Hoechst 33342 for 10 minutes. High-resolution fluorescence images were acquired either with a Zeiss LSM880 using ZEN2 software or with a Zeiss AxioObserver Z1 Apotome with AxioVision 4.8 software (Carl Zeiss, Jena, Germany) and processed with ZEN2 Software and Adobe Creative Suite 4 (Adobe Systems Incorporated, San Jose, CA).

All other images were acquired on the Leica TCS SPE laser scanning confocal microscope and processed using FIJI and Adobe Photoshop CS5.1 unless otherwise specified.

### Expression of GFP-tagged proteins in Zebrafish embryos

Capped mRNAs were synthesized using mMESSAGE mMACHINE SP6 transcription kit (Invitrogen Ambion). Embryos were injected at the 1-2 cell stage with 125-250pg of RNA.

### *Xenopus* ENKUR-GFP expression, morpholino knockdown and imaging

*X. laevis* embryo manipulations and injections were carried out using standard protocols. For ENKUR-GFP, capped mRNA was synthesized using mMESSAGE mMACHINE SP6 transcription kit (Invitrogen Ambion). A morpholino antisense oligonucleotide (MO) against *Enkurin* was designed to block splicing (Gene Tools). Its sequence was 5′- AATGACTATCCACTTACTTTCAGCC - 3′. mRNA and MOs were injected into two ventral blastomeres or two dorsal blastomeres at the 4-cell stage to target the epidermis or the gastrocoel roof plate (GRP), respectively. Each mRNA or MO was used at the following dosages: GFPEnkurin (60 pg), centrin4-BFP (40 pg), memRFP (50 pg) and Enkurin MO (20 ng). To verify the efficiency of *Enkurin* MO, MO was injected into the all cells at 4-cell stage and then RT-PCR was performed as described (Toriyama et al., 2016) (see table S9 for primers).

Confocal images were captured with LSM700 inverted confocal microscope (Carl Zeiss) with a Plan-APOCHROMAT 63×/1.4 oil immersion objective. Bright field images were captured on a Zeiss Axio Zoom. V16 stereo microscope with Carl Zeiss Axiocam HRc color microscope camera.

### Immunoblotting

Tissue was lysed using RIPA buffer (50mM Tris, pH 7.4, 150mM NaCl, 1% NP-40, 0.5% sodium deoxycholate) supplemented with protease inhibitors (Calbiochem, Billerica, MA). Protein samples were separated on 4–15% gradient TGX precast gels (Bio-Rad, Hercules, CA), transferred to nitrocellulose membrane (Whattman, Pittsburgh, PA). Membranes were blocked and antibodies were diluted in 5% milk in TBST (50mM Tris, pH 7.6, 150mM NaCl, 0.1% Tween 20) and analyzed using ECL Lightening Plus (Perkin–Elmer, Waltham, MA).

### Antibodies

Primary antibodies used were anti-β-actin (60008-I-Ig, ProteinTech, Rosemont, IL and ab8227, Abcam, Cambridge, MA), anti-TUB^ac^ (clone 6-11B-1, Sigma-Aldrich, St. Louis, IL), anti-b-tubulin (E7, Developmental Studies Hybridoma Bank, Iowa City, Iowa), anti-GFP (ab290, Abcam and sc-9996, Santa Cruz Biotechnology, Inc., Dallas, Texas), anti-ARL13B (clone N295B/66, Neuromab, Davis, CA and 17711-1-AP, ProteinTech), anti-ENKUR (HPA037593, Atlas Antibodies, Stockholm, Sweden), anti-ENKUR for SIM imaging (gift from H. Floreman, unpublished), anti-CCDC39 (HPA035364, Sigma), anti-CCDC151 (HPA044184, Atlas Antibodies), anti-DNAH5 (previously reported (Omran et al., 2008a)), anti-GAS8 (HPA041311, Atlas Antibodies), anti-DNAH11(previously described (Dougherty et al., 2016)), anti-RSPH9 (HPA031703, Sigma), anti-DNALI1 (HPA028305, Atlas Antibodies), anti-CCDC11 (HPA041069, Atlas Antibodies).

All secondary antibodies for immunofluorescent staining were Alexa-conjugated (Invitrogen). Hoechst 33342 was obtained from Invitrogen or Sigma-Aldrich. DAPI was obtained from Sigma-Aldrich. All horseradish peroxidase secondary antibodies for immunoblotting were obtained from Jackson ImmunoResearch Laboratories, Inc. (West Grove, PA).

### Quantitative RT-PCR

RNA was extract from cells using the RNeasy Mini or Micro Kit (Qiagen, Hilden, Germany) and cDNA was synthesized using iScript cDNA Synthesis Kit (BioRad). Quantitative PCR was performed using EXPRESS SYBR GreenER with Premixed ROX (Invitrogen) and the 7900HT Fast Real-Time PCR System (ThermoFisher Scientific). Relative expression was calculated using the delta-delta CT method (Livak and Schmittgen, 2001). For measuring expression of *Enkur,* the geometric mean of *Ubiquitin C* (*Ubc*) and *Hydroxymethylbilane synthase* (*Hmbs)* expression was used as the endogenous control for normalization. See Table S9 for primer sequences. The data were analyzed using Microsoft Excel and plotted using Prism7.

### Expression constructs and *in situ* probe plasmid generation

To construct the GFP-tagged expression plasmids for *S. purpuratus* ENKUR, ARRB1 and OPRM1L, gene specific cDNA was made from RNA of gastrula stage embryos using AccuScript (Agilent) or SuperScriptIII (Thermo Fisher Scientific), PCR amplified using primers containing the cutsites indicated below and TOPO-TA cloned into PCR2.1 (Invitrogen). The following restriction enzymes were used to insert cloned cDNA in the pCS2+8CeGFP (Gökirmak et al., 2012): for ENKUR - AscI and ClaI, for ARRB1 - AscI and PacI, OPRM1L - BamHI and ClaI. Unless otherwise indicated, the reverse primer for each gene was used to make gene specific cDNA. To construct the *L. variegatus* in situ probe, 480bp fragment of *L. variegatus Enkurin* synthesized using gBlocks Gene Fragments (IDT, San Diego, CA).

To generate the *X. laevis* ENKUR-GFP expression construct, full length of *Enkurin* cDNA was amplified from *X. laevis* RNA and cloned into pCS10R vector fused with N-terminal GFP using SalI and NotI. To generate the *X. laevis* in situ probe, the above plasmid was cut with SalI and synthesized using the T7 promoter. *Pitx2c* in situ probe, membrane-RFP (CAAX motif fused to RFP) and Centrin 4-BFP plasmids described in (Toriyama et al., 2016).

To create GFP fusions of human proteins to screen for ciliary localization in IMCD3s and *D. rerio* embryos human cDNA was cloned into pCS-eGFP-DEST using the Gateway system.

To construct the *S. rosetta* ENKUR-GFP expression plasmid, full-length cDNA (gift from Nicole King) was amplified, subcloned into pDONR221 and then cloned into pCS-eGFP-DES using the Gateway system.

To generated the mouse *Enkur* in situ probe, cDNA from *M. musculus* testis served as template for PCR for cloning a 598 bp fragment of *Enkur*. Via TOPO cloning, the fragment was ligated into the pCRII-TOPO vector (K4610, ThermoFisher Scientific,). The plasmid for the *Lefty2* in situ probe was a gift from R. Harland (Meno et al., 1997). The IMAGE clone ID 790229

The mouse *Enkur* shRNAs were expressed in pLKO.1 and were purchased from Sigma-Aldrich, MISSION shRNA collection (see Table S9 for clone numbers).

All primer and shRNA sequences are listed in Table S9.

### In situ hybridization

#### Mus musculus

For the mouse *Enkur* in situ probe, plasmid was digested with NotI for antisense probes and BamHI for sense control probes and purified using phenol and chloroform (see Table S9 for primers). Digested plasmid DNA served as template for *in vitro* transcription with Sp6 (antisense) or T7 (sense) RNA polymerases to generate digoxigenin (DIG) labeled RNA probes (Roche). For the *Lefty2* probe, plasmid was digested with EcoRI and *in vitro* transcribed using T7 RNA polymeras. For the *Cerl2* probe (IMAGE clone ID 790229), plasmid was digested with NotI and *in vitro* transcribed using T7 RNA polymerase.

Embryos were fixed in 4% PFA in PBS for at least 24 hours, transferred to methanol and stored at -80°C until use or, if in situ was performed within 2 weeks of dissection, embryos were stored in fixative at 4°C. If in methanol (Fig S5 only), embryos were rehydrated to PBST, bleached with 6% H_2_O_2_ for 1 hour, digested with 10μg/ml proteinase K (Roche) for 8 minutes and fixed with 4% PFA and 0.2% glutaraldehyde. The embryos were incubated in hybridization solution (50% formamide, 0.5% CHAPS, 0.2% Tween, 1.3x SSC, 5mM EDTA, 50μg/ml yeast t-RNA; 700U/ml Heparin) at 65°C for several hours. Hybridization with the DIG-labeled RNA probes was performed overnight at 65°C in hybridization solution. The following day, embryos were washed two times with fresh hybridization solution for 1hr each, transferred to MABT (100mM Maleic acid, pH7.5, 150 mM NaCl, 0.5%Tween) and washed 3 times with MABT for 15 minutes each. After incubating with blocking solution (2% Boehringer Blocking Reagent (Roche), 20% heat-treated sheep serum in MABT) for 1 hour the embryos were incubated overnight at 4°C with AP-anti-DIG antibody diluted in blocking solution. Following 4 washing steps in MABT for 1 hour each, and 1 wash overnight at 4°C the embryos were transferred to NTMT (100mM NaCl, 100 Tris, pH9.5, 50mM MgCl_2_, 0.1% Tween). Color was developed using NBT/BCIP or MB Purple Substrate (Roche). Embryos were imaged using a Zeiss Discovery.V12 stereomicroscope and AxioCam MRc camera or a Nikon Digital Sight DS-L3 camera mounted on a Nikon SMZ1000 stereomicroscope. Images were processed using creative suite (Adobe).

#### Xenopus

Whole mount in situ hybridization of *X. laevis* embryos was performed as described previously (Sive et al., 2000) using DIG-labeled single-stranded RNA probes against *Enkurin* and *Pitx2c* (see Table S9 for primers).

#### Sea urchin

Whole mount in situ hybridization of *Lytechinus variegatus* embryos was performed as described previously (Warner et al., 2016) using DIG-labeled single-stranded RNA probes against a *Enkur* (see Table S9 for primers).

#### Sea anemone

Whole mount in situ hybridization of *N. vectensis* embryos was performed as described previously (Bause et al., 2016) using DIG-labeled single-stranded RNA probes against *Enkurin* (see Table S9 for primers).

### High speed video microscopy of cilia of the mouse tracheal epithelium

Mouse trachea were dissected and imaged as described in (Francis and Lo, 2013). Video were captured on a Nikon Ti with PFS3 inverted microscope using a Andor Zyla 5.2 camera.

### Human subject phenotyping

We obtained signed and informed consent from patients fulfilling all PCD diagnostic criteria as well as their family members using protocols approved by the Institutional Ethics Review Board of the University of Muenster (Germany). PCD phenotype was characterized by standard clinical diagnostic criteria and documentation of typical symptoms such as neonatal respiratory distress and signs of chronic oto-sinu-pulmonary infections with development of bronchiectasis. Clinical diagnosis included nasal nitric oxide (NO) measurement, medical imaging (X-Ray), high speed video microscopy (HVMA), transmission electron microscopy (TEM) as well as immunofluorescence analysis (IF) to analyze ciliary structure and function. The value of nasal NO was measured while performance of an exhalation-against-resistance maneuver, as previously described (Olbrich et al., 2012). For HVMA, cells obtained by nasal brush biopsy were spread on glass slides and immediately ciliary beating was analyzed using the SAVA imaging analysis system (Ammons Engineering, Mt. Morris, MI). Equipment and settings were previously described (Olbrich et al., 2015).

### Haplotype analysis, genome-wide SNP mapping, whole exome sequencing and Sanger sequencing

Haplotype analysis and genome-wide single nucleotide polymorphism (SNP) mapping were performed as previously described (Olbrich et al., 2012). Exome sequencing of genomic DNA was performed at the Cologne Center for Genomics (CCG). For enrichment, the NimbleGen SeqCap EZ Human Exome Library v2.0 was used. Enriched preparations were sequenced with the HiSeq2000 platform (Illumina) as paired end 2x 100 bp reads. Sequencing reads that passed quality filtering were mapped to the reference genome sequence (hg19). Variants were analyzed using the varbank software (previously described (Olbrich et al., 2015)). In addition to the c224-1delG mutation described above (see Results), two other SNPs were detected in *ENKUR* that segregated with the disease phenotype (Table S7). These SNPs have allele frequencies of 3% and 5%, indicating that they do not cause rare phenotypes like situs inversus. PCR amplification of genomic fragments and Sanger sequencing was performed as previously described (Tarkar et al., 2013) to analyze all *ENKUR* exons including the exon intron boundaries (See Table S9 for primers).

### Mouse strains

*Enkur* mutant mice were obtained from H. Floreman and are described in (MS not published yet). Briefly, a targeting vector was used to delete a portion of exon 2 and insert a strong splice acceptor followed by a neomycin resistance cassette. As exons 1 and 3 are in different reading frames, any splicing around the cassette will produce a frameshift. Mice were maintained on a mixed C57BL/6J-129 genetic background. The Institutional Animal Care and Use Committee at the University of California, San Francisco approved all protocols for the *Enkur* mouse line protocols.

A C57BL/6J male mouse was used for the isolation of RNA from testis for generation of the *Enkur* in situ hybridization plasmid. CD-1 wild type embryos were used for *Enkur* in situ hybridization (Figure S5F). All animal experiments with the CD-1 mice complied with ethical regulations and were approved by local government authorities (Landesamt für Natur, Umwelt und Verbraucherschutz Nordrhein-Westfalen, Germany; AZ 84-02.05.20.12.164, AZ 84- 02.05.20.12.163, AZ 84-02.05.50.15.025 and AZ 84-02.05.50.15.012).

## Author Contributions

Conceptualization, M.A.S. and J.F.R.

Methodology, M.A.S., H.O., J.J., N.K. F.R., J.B.W. and J.F.R.

Investigation, M.A.S., T.M., J.J., C.L., S.P.C., G.G.3^rd^, H.B., G.D., P.P., C.W.

Resources, H.O., N.K. F.R., J.B.W. and J.F.R.

Writing – Original Draft, M.A.S. and J.F.R.

Writing – Review & Editing, M.A.S., J.F.R., T.M., and F.R.

## Acknowledgments

We thank Fred Wilt and Christopher Killian for providing sea urchin embryos for initial experiments and for guidance with sea urchin husbandry, Stefan Materna for sea urchin husbandry assistance, Nicole King for providing the choanoflagellate culture line and for experimental advice, Daniel Richter, Tera Levin and Pawel Burkhardt for counsel on choanoflagellate culture and experiments, Hannah Elzinga for cloning a choanoflagellate gene, Harvey Florman for sharing the *ENKUR* mutant mice, David R. McClay and Esther Miranda for *L. variegatus* embryos and the UCSF Nikon Imaging Center for providing microscopes and imaging assistance. We thank the PCD individuals and their families for participating in this study and acknowledge the German patient support group “Kartagener Syndrom und Primaere Ciliaere Dyskinesie e.V.” We thank H. Olbrich and N.T. Loges for fruitful discussions and M. Herting, B. Lechtape, A. Robbers, D. Ernst, L. Overkamp, K. Wohlgemuth and F.J. Seesing for excellent technical assistance, G. Nürnberg and P. Nürnberg for homozygosity mapping data, and the Exome Aggregation Consortium (ExAC) for creating their exome variant database.

This work was supported by grants from the NIH (AR054396 and GM095941) and from the Burroughs Wellcome Fund, the Packard Foundation, and the Sandler Family Supporting Foundation to J.F.R. Work in the lab of H.O. was supported by the Deutsche Forschungsgemeinschaft (OL/450-1 (H. Olbrich) and OM 6/4, OM 6/7, and OM6/8 (H. Omran), Interdisziplinären Zentrum für Klinische Forschung Muenster grant Om2/009/12 and Om2/015/16 (H. Omran), European Commission grant FP7/2007–2013 grant agreement (GA) 262055 (H. Omran) as a Transnational Access project of the European Sequencing and Genotyping Infrastructure, EU-FP7 programs SYSCILIA GA 241955 and BESTCILIA GA 305404 (H. Omran), the Eva Luise und Horst Köhler Stiftung and Kindness for Kids.

